# A database of accurate electrophoretic migration patterns for human proteins in cell lines

**DOI:** 10.1101/2022.06.22.496709

**Authors:** Roman Mylonas, Alexandra Potts, Patrice Waridel, Jachen Barblan, Maria del Carmen Conde Rubio, Christian Widmann, Manfredo Quadroni

**Affiliations:** Protein Analysis Facility, Faculty of Biology and Medicine, University of Lausanne, Lausanne, Switzerland; Department of Biomedical Sciences, Faculty of biology and Medicine, University of Lausanne, Lausanne, Switzerland

**Keywords:** proteins, molecular weight, electrophoresis, SDS-PAGE, mass spectrometry, internal calibration, differential splicing, post-translational modifications

## Abstract

Native molecular weight (MW) is one of the defining features of proteins. Denaturing gel electrophoresis (SDS-PAGE) is a very popular technique for separating proteins and determining their MW. Coupled with antibody-based detection, SDS-PAGE is widely applied for protein identification and quantitation. Yet, electrophoresis is poorly reproducible and the MWs obtained are often inaccurate. This hampers antibody validation and negatively impacts the reliability of western blot data, resulting worldwide in a considerable waste of reagents and labour. To alleviate these problems there is a need to establish a database of reference MWs measured by SDS-PAGE. Using mass spectrometry as an orthogonal detection method, we acquired electrophoretic migration patterns for approximately 10’000 human proteins in five commonly used cell lines. We applied a robust internal calibration of migration to determine accurate and reproducible molecular weights. This in turn allows merging replicates to increase accuracy, but also enables comparing different cell lines. Mining of the data obtained highlights structural factors that affect migration of distinct classes of proteins. We also show that the information produced recapitulates known post-translational modifications and differential splicing and can be used to formulate hypotheses on new or poorly known processing events. The full information is freely accessible as a web resource through a user friendly graphical interface (https://pumba.dcsr.unil.ch/). We anticipate that this database will be useful to investigators worldwide for troubleshooting western blot experiments, but could also contribute to the characterization of human proteoforms.

## Introduction

The ability to reliably resolve, identify and quantify individual proteins is essential in both fundamental biology and biomedical research. For separation and profiling of complex protein mixtures, sodium dodecyl sulfate (SDS)-based electrophoresis (SDS-PAGE) has played a central role since its establishment by J. Maizel and U. Laemmli^1,2^. For protein identification, mass spectrometry (MS) can identify proteins based on their most integral property, the amino acid sequence. However, due to its cost and complexity, MS cannot be implemented in all labs. Thus in a majority of laboratories, specific protein detection is performed by western blot (WB), which combines MW-based separation with recognition of the protein of interest by a specific antibody (Ab). WB has grown in popularity thanks to its low cost, ease of implementation and, at its best, exquisite sensitivity and specificity. However, WB is also often plagued by artefacts and technical issues at the separation and membrane transfer steps, in addition to major issues due to the reliability of the Ab’s used. Detection antibodies are essential tools in fundamental biology, biomedicine and medical diagnostics. And yet, they have also been singled out as major contributors to the poor reproducibility of scientific publications ^3–8^. The reasons for this situation, as well as its considerable scientific and economic costs, have been previously discussed ^9^. In practice, the problem can be reduced to two underlying issues, namely i) the often insufficient characterization and validation of commercial Ab’s and ii) the wide range of biological systems and techniques in which Ab’s are used. The combination of these two variables recurrently creates problems and ambiguities whenever a new Ab must be validated for a specific application in a given class of samples. While the Ab market has been estimated at around 800 million US$ yearly ^9^, the cost in manpower dedicated annually to troubleshooting antibody-related problems is impossible to quantify but is likely much bigger than the cost of the reagents themselves.

For WB, an ideal Ab should be specific and only generate a single or several band(s) at the known molecular weight(s) of the target protein species. The latter is a key parameter which can be known experimentally for the mature protein, but most often is simply inferred from the gene- or mRNA-derived sequence. In everyday practice a number of issues often arise that can make validation of WB results difficult, namely i) a WB band is observed but its apparent MW on gel does not correspond to the expected MW calculated from the sequence ii) a WB band is observed but its apparent MW does not correspond to the one specified in the Ab data sheet, which in turn is often determined approximatively and in a different type of sample, iii) several bands are observed, of which only one or none corresponds to the expected MW and iv) the band(s) observed are different in number and MW in different tissues or samples.

These numerous issues have different origins, in part related to Ab properties as described above and in part due to the WB technique, which is complex, manual and poorly reproducible. Also, despite the advent of CRISPR/Cas9 techniques, a negative control not expressing the target protein is not always available. The complexity and plasticity of protein processing, especially in eukaryotic cells, in which differential mRNA splicing and post-translational modifications (PTMs) are widespread are additional confounding factors. These phenomena can alter dramatically protein MW and affect electrophoretic migration and the pattern observed in a particular sample. Thus in many cases an investigator performing a WB for the first time does not know, beyond the rather simplistic assumption of “one band at the right place”, what pattern (number of bands, apparent MW) should be expected for the protein of interest in a specific sample. It follows from these considerations that one of the main limitations at the moment is the lack of accurate and reliable information on the actual migration of proteins in SDS-PAGE. Such knowledge, if available, would facilitate the validation (or dismissal) of antibody reagents as well as WB results in general. Ideally, such reference data on gel migration should be obtained in a manner fully orthogonal to Ab-based detection and thus should be created from MS-based identifications.

Gel electrophoresis has been widely applied as a protein separation method before MS analysis in so-called geLC-MS workflows^10^. In such experiments, proteins are separated by classical SDS-PAGE, followed by cutting of desired gel regions and in-gel digestion with (mostly) trypsin. Peptide fragments are recovered in solution and submitted to nanoLC-MSMS for protein identification. GeLC-MS approaches are robust and compatible with many types of samples and can yield extensive pre-fractionation to reduce sample complexity. In recent years, with increased speed and resolution of MS systems, the needs for sample prefractionation decreased and, at least for total proteome studies, other in-solution workflows were often preferred, which offer higher throughput and sample recovery. Extensive gel fractionation has continued to be applied for projects in which a correlation of identification with protein MW is essential, e.g. for mapping proteolytic events. We have used extensive gel fractionation, coupled with SILAC labelling, to identify the cellular substrates of the NS3-4A Protease from Hepatitis C Virus^11^ as well as the effects of protease inhibition^12^. A similar approach was used by others to map cleavage events linked to apoptosis^13,14^. In these studies, the goal was to identify shifts in protein migration induced by cleavage or PTM’s in general. However, it is easy to see that extensive gel-based fractionation into 45-50 gel slices, when coupled with MS-based identification of proteins in each slice, does also produce detailed maps of protein presence as a function of position in the gel, as reported ^15^. Since data is obtained for >4000 proteins in a single experiment, the information could be exploited for evaluating the specificity of a large number of antibodies. Indeed, the Human Protein Atlas team has chosen this strategy as one of the pillars used to evaluate more than 6000 Ab produced by the consortium ^16^.

We decided to use a similar approach but with a different scope, i.e. to build an easily accessible, user-friendly reference database of SDS-PAGE migration patterns for human proteins. We analysed with an extended geLC-MS workflow extracts of 5 widely used human cell lines and generated detailed protein ID and quantitative data as a function of gel migration position. Crucially, we have developed a dedicated data analysis pipeline to determine accurate, internally-referenced gel MW values without using external markers. We then used this information to align and average three replicate runs for each line, thus increasing accuracy and robustness of the data. MW calibration also allows to carry out accurate comparisons of different cell lines. The information is available via a freely accessible web resource (https://pumba.dcsr.unil.ch/) with several options for graphical output including peptide coverage maps. We also show that the database can be mined to derive novel observations on so far unknown protein processing events. We argue that this database will be broadly useful to investigators using WB techniques in order to evaluate their results for proteins of interest. Furthermore, when integrated into existing knowledgebases, the migration patterns we determined could be useful for elucidating the occurrence of human proteoforms.

## Materials and Methods

### Reagents

Unless specified otherwise, all reagents were of analytical grade purchased from Merck-Sigma (Buchs, Switzerland).

### Sample preparation

U2OS and HCT-116 cells were purchased from ATCC as described^17^. All other cell lines were purchased from ECACC (European Collection of Authenticated Cell Cultures) through a local distributor (Millipore-Sigma). Adherent cells were grown to half dish confluence, after which the medium was removed by aspiration and layers of cells were washed with PBS at 4°C. Cells were then mechanically scraped in FASP buffer (4% SDS 100mM DTT, 100mM Tris pH 7.5)^18^ and the resulting suspension was immediately heated at 95°C for 5 min, sonicated with a tip sonicator for 3×20s and centrifuged at 13’000 xg for 10 min. Protein concentration in the supernatant was determined with the tryptophan fluorescence method^19^. Suspension cells grown to a density of 5 × 10^5^ cells/ml were diluted 5x with ice-cold PBS and recovered by centrifugation at 1’200 x g for 2 min. After washing two more times with excess ice-cold PBS, cells were lysed in FASP buffer as described for adherent cells. Samples for U2OS and HCT-116 cells were prepared as SILAC samples. While the light (unlabelled) sample was treated with 4 μM cisplatin for 48 h, the heavy isotope-labelled sample was control cells treated with vehicle only (DMSO). Only the intensities of the heavy control channel were used for database construction.

### Electrophoresis and protein digestion

For protein separation in the case of SILAC-labelled samples, Heavy and Light labelled lysates were mixed at a quantitative ratio of 1:1. For all cell lines, a total of 120 μg of protein from lysates was migrated on a pre-cast 4-12% Novex NuPAGE SDS Mini Gel (Invitrogen/Thermo Fisher) using the Bis/Tris buffer system (product number NP0326BOX) with MOPS running buffer, according to instruction from the manufacturer. Gels were stained overnight with colloidal Coomassie blue^20^. Entire lanes were cut into 45-47 identical gel slices using a gridcutter tool (MEE1-5-50, The Gel Company, San Francisco, CA, USA) (Supplementary Fig. S1). In-gel proteolytic cleavage with sequencing grade trypsin (Promega) was performed robotically, according to a described protocol ^21^. The supernatant from the digestion was concentrated by evaporation and redissolved in 30 μl 0.05% trifluoroacetic acid (TFA) in 2% acetonitrile for liquid chromatography-tandem mass spectrometry. Samples were analyzed on a Q-Exactive Plus mass spectrometer (Thermo Fisher Scientific) interfaced *via* a nanospray source to an UltimateRSLC 3000 nanoHPLC system (Thermo Fisher Scientific). Peptides were separated on a reversed-phase Acclaim PepMap nanocolumn (75 μm inner diameter x 50 cm, 2 μm particle size, 100 Å pore size; Dionex) or a reversed-phase, custom-packed nanocolumn (75 μm ID x 40cm, 1.8 μm particles, Reprosil Pur, Dr Maisch) with a 4-76% acetonitrile gradient in 0.1% formic acid (total time 65 min). Full MS survey scans were performed at 70’000 resolution. In data-dependent acquisition controlled by Xcalibur software (Thermo Fisher), the 10 most intense multiply charged precursor ions detected in the full MS survey scan were selected for collision-induced dissociation (HCD, normalized collision energy NCE=27%) and analysed in the orbitrap at 17’500 resolution. Selected precursors were then excluded from selection during 60 s.

### MS data analysis

GeLC-MS data were analyzed with MaxQuant version 1.6.14.0, using the Andromeda search engine for identification ^22,23^. Database searches were performed on the reviewed set of canonical human sequences of the UniProtKB/SWISSPROT database, downloaded on September 16^th^, 2020 containing 20’371 sequences and supplemented with common contaminants. Search parameters were trypsin specificity, two possible missed cleavages and mass error tolerances of 7 ppm for the precursor and 20 ppm for tandem mass spectra after recalibration. The iodoacetamide derivative of cysteine was specified as a fixed modification, and oxidation of methionine and protein N-terminal acetylation were specified as variable modifications. Peptide and protein identifications were filtered at 1% false discovery rate established against a reversed database, according to default MaxQuant parameters. A minimum of one unique peptide was necessary to discriminate sequences which shared peptides. Sets of protein sequences which could not be discriminated based on identified peptides were listed together as protein groups. All gel slices in a run were analysed as separate experiments by MaxQuant, with the “match between run” option activated applied between adjacent slices. Data for the three replicates for each cell line were analysed together in a MaxQuant run, followed by processing and import into the database. Thus, protein inference and definition of protein groups was done at the level of each cell line.

The mass spectrometry proteomics data have been deposited to the ProteomeXchange Consortium ^24^ via the PRIDE partner repository with the dataset identifier PXD026750. The dataset is available on https://www.ebi.ac.uk/pride/archive with the following information : username , password: g6TBDKtI.

### Database design and data import

MaxQuant result files (proteinGroups.txt and peptides.txt) for each cell line were transformed and imported into a document database (MongoDB). The corresponding code and database design are available on GitHub (https://github.com/UNIL-PAF/pumba-backend). As a first step, contaminants were removed using the corresponding MaxQuant annotation (column “Potential contaminant”) and an additional list of environmental contaminants generated in the lab. Then the result files were parsed and stored in 3 different entities in the database (MongoDB): <u>datasets</u>, <u>proteins</u> and <u>sequence</u> (Supplementary Figure S2A). A dataset in the context of Pumba consists of a single replicate from a cell line. Cell line name, replicate name, species, slice-to-mass fits (described later) and a normalization factor (calculated from the sum of all protein group intensities in the dataset) are stored in this table. Next, every protein group for every dataset is stored as a separate entry in the proteins table. For every entry the majority protein ID’s (proteins that have at least half of the peptides that the leading protein has), protein names and gene names and the corresponding peptides are imported. For every peptide, the majority protein ID’s it belongs to, the peptide sequence, the amino acid preceding and following the sequence, the theoretical mass and Andromeda score are recorded, together with the information whether it is a razor peptide or if it is unique by group. Finally the raw intensity of the peptide in each slice is recorded. To create the sequence table, the same FASTA files that were used for the MaxQuant identification were parsed and imported. For every protein entry the corresponding AC, protein and gene names, sequence and sequence length are stored. The theoretical mass for each protein was calculated from its sequence.

### Global slice-to-mass fitting

The following steps were done in R and implemented in a R package developed in house (https://github.com/UNIL-PAF/pumbaR). To estimate the molecular weight for each gel slice, the MaxQuant proteinGroups.txt result files were imported as a table. First all contaminants and reverse hit proteins were removed and only the first protein match from each protein group was used. For each protein and slice the intensity information was used (MaxQuant “Intensity” column and for SILAC labelled cells (U2OS and HCT-116) the “Intensity H” column). Protein intensities from a few gel slices with low content and many contaminants (such as at the lower and upper boundaries of gels), thus showing a very atypical behaviour, were set to 0 so they were not used in any of the following steps. The molecular mass of each slice was then estimated by fitting a third degree polynomial function to a plot of theoretical protein masses against its slice position in the gel (Figure 2A, upper panel). The protein intensities in each slice were used as weights for the fitting. To further improve the fit, only a subset of all proteins was used. First, proteins that appeared in more than 30% of the slices (probably contaminants e.g. keratins) were removed. Second, proteins with intensity below 50% of the max intensity were removed. Third, proteins from regions in the plot with only few data points were removed. This was done by laying a grid of 500 × 500 over the plot and only keeping the data in cells with more than 10 proteins. In the end around 30% of all data points were used (e.g. for replicate 12’019 from cell line HEK293 only 13’216 data points from 7’776 proteins were used out of a total of 45’744 points for 8’374 proteins) (Figure 2A, lower panel). For every dataset a resulting table with estimated molecular weights for each gel slice was obtained and stored in the database.

**Figure 1.**
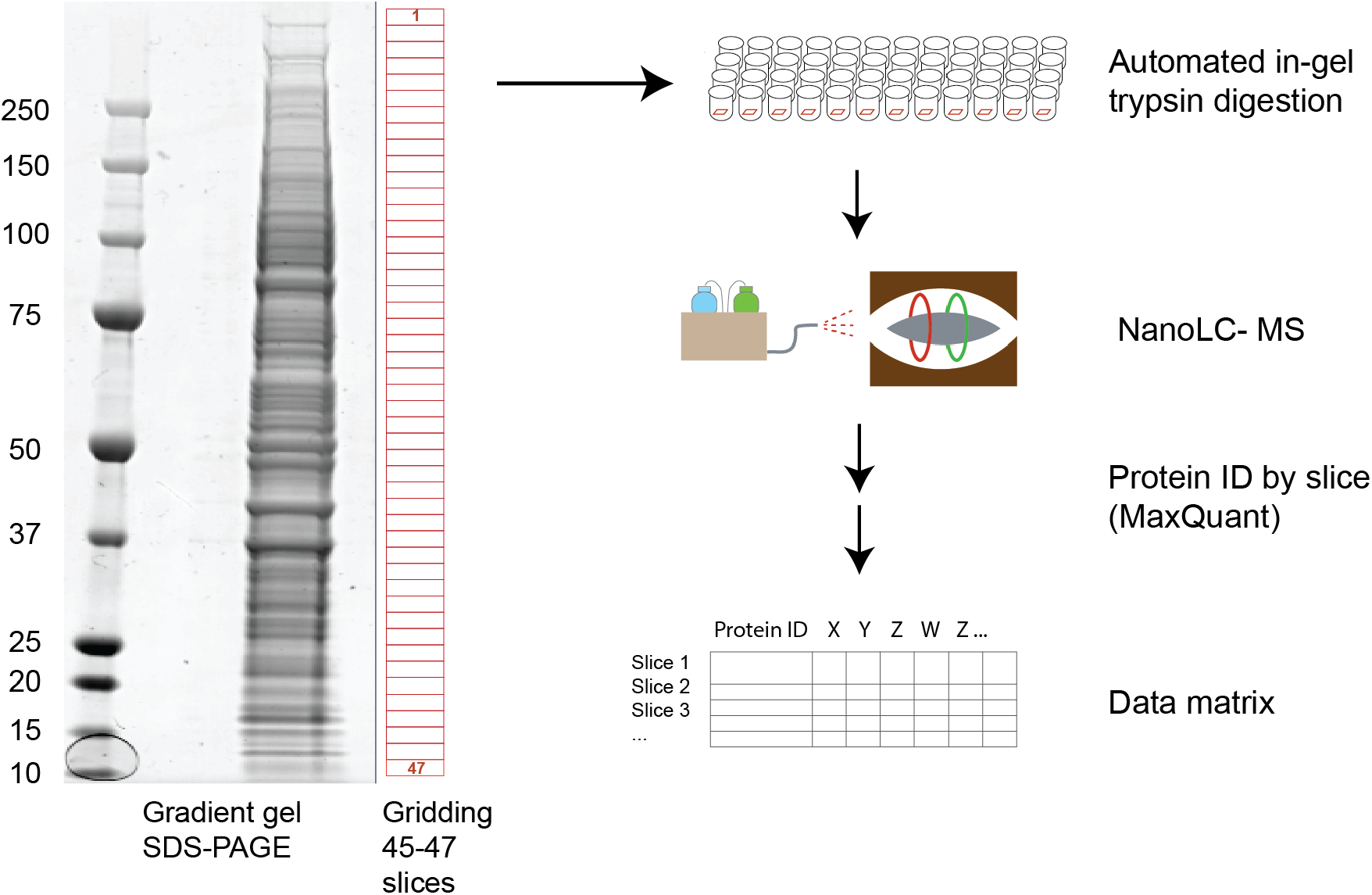
Workflow for protein separation and data acquisition. Cell extracts were separated on a standard precast gel system, stained and excised into 45-47 slices. Robotized in-gel digestion and LC-MS/MS analysis generated protein identifications and intensity data for all gel slices.

**Figure 2.**
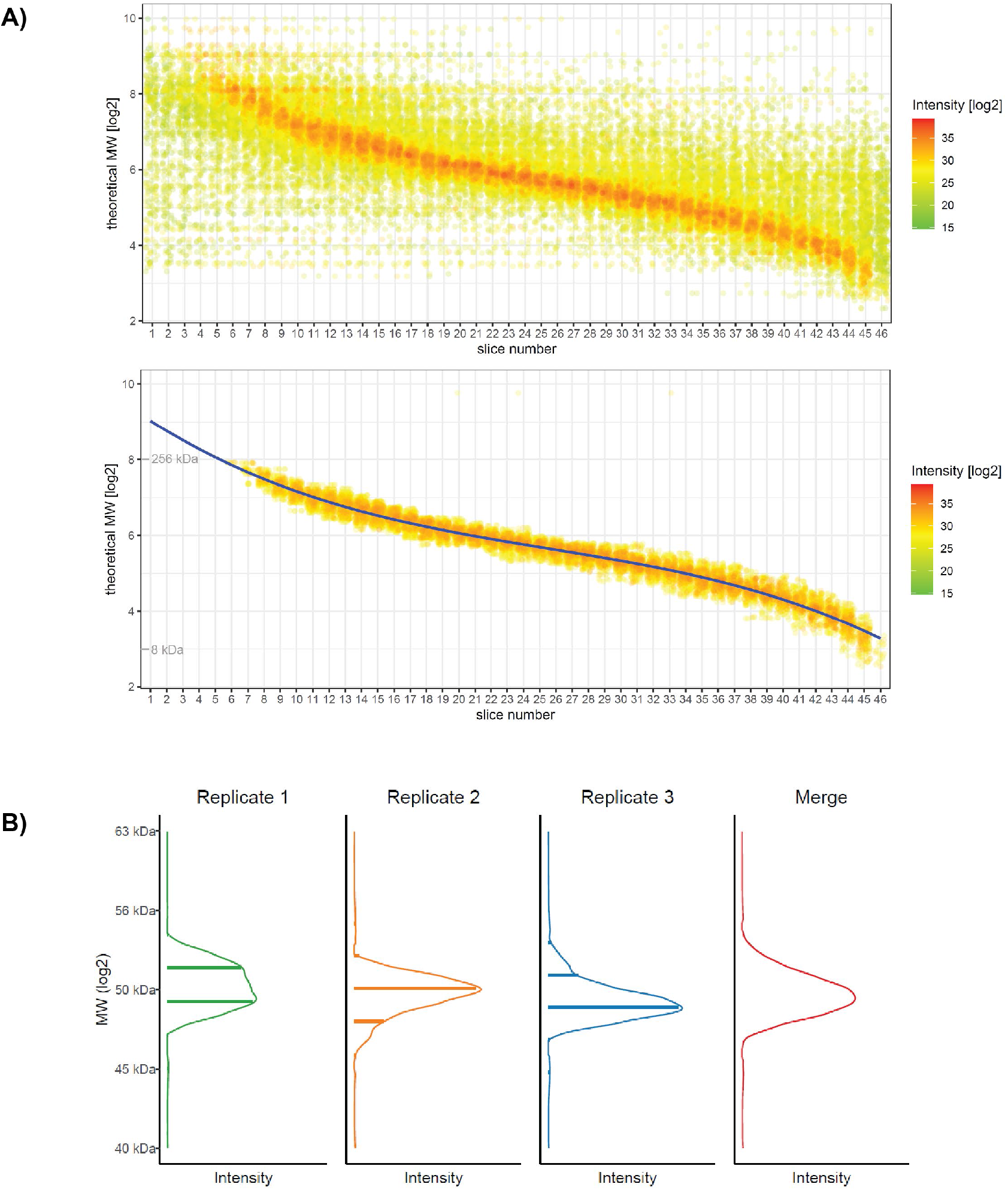
Raw data, MW distribution and fitting. **A)** Plot of theoretical (calculated from the sequence) MW for identified proteins vs. identification in individual gel slices. Every point is a protein identification instance. Raw data (upper panel) and filtered data (lower panel) are shown for a representative replicate (HEK293 cells, replicate 3). A third-degree polynomial function is fitted on the data after filtering to keep the strongest signals (lower panel). The polynomial function is then used to determine slice center mass, allowing alignment of replicates. **B)** Interpolation, merging of replicates and smoothing create the final profile. For details on the procedure see Supplementary Figure S2.

### Querying the database to retrieve and display data for a protein

To query the database, a user has to provide a query protein AC or gene name and the list of datasets to be loaded. All protein groups and datasets which do contain the given protein AC or gene name are loaded from the database. All information from the protein groups and all peptides corresponding to the corresponding protein AC are retrieved. The total protein intensity for the selected protein is computed by summing up all individual peptide intensities. The position of each peptide within the protein sequence is recomputed on every user request based on the sequence queried. The protein and peptide intensities of each dataset are then normalized using the normalization factor previously calculated from the total intensity in the dataset. In case the query is <u>not</u> the leading protein in a protein group, a flag is added in the data display (“lanes” view, when replicates are shown) to highlight the fact that there was an overall better candidate sequence identified by a higher number of peptides and that therefore the presence of the queried entry cannot be conclusively established in that dataset.

### Interpolation of intensities across gel slices

A merge of all the replicates for every cell line is calculated for the query protein using a method implemented in Scala (https://github.com/UNIL-PAF/pumba-backend)(Figure 2B, Supplementary Figure S2B). For this, the normalized protein intensities for the selected protein groups are retrieved by cell line. Then, for each dataset within a cell line group, a table of masses and intensities is created and every slice is divided into 100 microslices along the MW coordinate, with the intensity spread uniformly among them. Afterwards the microslices from the different datasets are aligned, matched by MW within a certain tolerance and the average of the intensities is taken. If only one dataset is selected for display, the microslices are taken as they are. A LOESS curve is fitted through the averaged intensities, and intensity and mass information of the fit are retained (around 5000 data points). The result is one curve (intensity *vs*. MW) per cell line, associated with the database entry that was queried.

### Visualization in the web interface

All normalized protein and peptide information together with the merges, are retrieved, combined with the sequence information and are subsequently used by the web interface to visualize the different plots (https://github.com/UNIL-PAF/pumba-frontend). In this version of the database, splice variants were not considered, due to the difficulty to assigning intensities in case of presence of two or more variants with extensive shared sequences. However, the predicted MW of UNIPROT-annotated splice variants can be visualized in the “lanes” view of the GUI.

### Processing of information extracted from the database for plotting and analysis

For QC and analysis of content, a raw table was exported from the database containing a large set of features (Supplementary table ST2, see table for description of columns). To create this table, data was extracted from the database by accessing Pumba as an API. The R scripts used for the table creation can be found under (https://github.com/UNIL-PAF/pumba-datamining). Values for pI and hydrophobicity were calculated in R with the EMBOSS and Hopp-Woods methods, respectively. Further information added during export was parsed from UNIPROT and included all annotations except GO, KEGG, Interpro, Corum, UNIPROT keywords. The latter were added using the Perseus software^25^ using data in https://annotations.perseus-framework.org. Perseus was also used for all subsequent steps, i.e. calculate numerical Venn diagrams, correlation coefficients (Pearson’s correlation), CVs and other general statistics. The initial main table exported from the database (Supplementary Table ST2) lists 10’187 proteins. For further analysis, the table was filtered to retain only proteins present in all replicates of all cell lines with a highest peak that contains at least 70% of the total intensity in that replicate (Supplementary Table ST4, 2’688 proteins). A subset of table ST4 was prepared, containing only the averaged cell line “highest peak” columns. The FMD (Fractional Mass Deviation) was calculated as (xxx.highest.peak.mass-theo.mass)/theo.mass from interpolated, averaged values (Supplementary Table ST5 (2’688 proteins)). 1D annotation enrichment was calculated with Perseus ^26^ based on sorting by the FMD values for each of the 5 cell lines, with Benjamini-Hochberg FDR correction of the enrichment with threshold at q-value<0.01. For further plotting and presentation, only annotation terms that were significant in at least 4 out of 5 cell lines were considered.

## Results

### Data acquisition

To build the database we analysed lysates of HeLa, HEK293, Jurkat, U2OS and HCT-116 cell lines. These lines were chosen because of their widespread use in research lab, and to represent different cell types. Total extracts were prepared under strong solubilizing conditions (4% SDS) to ensure comprehensive extraction of cell compartments. For adherent cells, lysis was done *in situ* on plate to avoid artefacts linked to cell detachment by trypsinisation. Gel separation was carried out with pre-cast, commercially available gradient gels to ensure reproducibility and high resolution over the largest possible range of MW (Figure 1). We used enlarged gel wells to increase loading and we only excised the portion of each lane with minimal streaks and border effects. The number of slices taken was between 45 and 47 (Supplementary Fig.S1, Supplementary table ST1). Replicates for cell lines corresponded to independently started cell cultures, sometimes prepared months apart (Table ST1). After in-gel trypsin digestion, proteins were identified by high resolution LC-MS/MS. MS data were analysed with MaxQuant^23^, whereby each gel slice was defined as a separate experiment. This allows monitoring the profile (signal intensity) of each identified protein, across all positions in the gel. The number of protein groups identified by MaxQuant ranged from approximately 5200 (HCT-116 cells) to 8400 (HEK-293 cells) per replicate (one full lane), filtered with standard MaxQuant parameters (1% peptide and protein FDR) at minimum 1 peptide (Supplementary Table ST1). U2OS and HCT-116 cells had lower numbers of IDs, due to their SILAC labelling format that increases spectral complexity.

### MW fitting, alignment and averaging of data from different samples

Our general goal was to determine accurate, robust values of apparent protein MW as determined by electrophoresis. Since proteins can be present as many species with potentially distinct apparent molecular weights, for the sake of clarity we will use the term “highest peak MW” to describe the molecular weight of the main, quantitatively dominant species detected. SDS-PAGE is not considered a highly reproducible technique and can be influenced by many environmental factors and variables. We thus reasoned that reference values should be obtained by averaging independent replicates. Key to the development of the database was thus the ability to determine MWs accurately and in a robust manner from a gel migration and to align results from any geLC-MS runs for averaging among replicates or comparison among cell lines. A plot of the observed raw protein intensity signals as a function of gel slice number and the logarithm of the theoretical (calculated from the sequence) MW (Figure 2A) showed, as expected, a sigmoidal correlation with a central linear domain. Rather than rely on a few external markers for calibrating the migration curve, we therefore exploited the wealth of information provided by thousands of MS identifications to calculate an internally-referenced calibration curve. This was based on the key assumption that a majority of proteins in the measurable proteome have a molecular weight that is close to the one predicted from the sequence and migrate approximately as expected from their MW. Filtering the data to retain only identifications with a higher number of peptide-spectrum matches (and thus stronger signal intensity) corroborated this assumption and greatly reduced the spread of the distribution (Figure 2A,lower panel). Polynomial fitting of the filtered dataset thus allowed calculation of an average MW value for each slice in each gel separation. Overall plots of protein distributions as a function of gel migration were highly similar across replicates and cell lines and fitting could be performed in all cases without major difficulties (data not shown).

The next step was the alignment and averaging of replicate runs. The data has a discrete structure, made of protein identifications and intensities linked to the theoretical centre of each gel slice in which they were detected, which in turn is a portion (bin) of the MW range. We developed a pipeline that interpolates normalized intensities in and across gel slices, then aligns (based on the fitted MW) and averages replicate runs to obtain the final profile for the cell line considered (Figure 2B, for details see Supplementary Figure S2B). The raw intensity values in each gel slice for all replicates are retained and can be visualized at any time in the database interface for verification. Local and global maxima of the obtained intensity curve were detected and integrated to yield quantified peaks that were then used for quality control and data mining. Overall, the separation for the gel/buffer system used appeared to fit an ideal behaviour in the range from 300 kDa down to 10 kDa. Very few proteins were detected beyond these boundaries.

### Database content and overall patterns observed

Initial tests of the workflow using technical duplicates prepared using the same sample showed highly reproducible results (data not shown). We thus proceeded to collect the datasets for all cell lines, fit and obtain slice molecular weights and populate the database. Database content for the human cell lines was then processed to extract essential metrics in a table format for inspection and quality control. All intensity peaks that were determined on interpolated data are reported in Supplementary Table ST2, with a dedicated column for the apex of the most intense peak and its integrated value. Position and intensity of such “highest peak”, which would represent the main band in a conventional gel or western detection, were analysed more in depth to determine its contribution relative to the total signal of the protein and its position relative to the theoretical, expected MW. Table ST2 summarizing the database contains data for 10’187 proteins. Of these, 3’535 had values in all replicates of all the 5 human cell lines. Numbers of proteins with values in the database for individual cell lines were in general very close (+/-2%) to the numbers of protein groups identified by MaxQuant in the same line, with slight differences due to the inference process used to align results from different lines. The percentage of proteins identified by a single peptide match varied from 3.3% (HEK 12019 sample) to 14% (U2OS 9053 sample), depending on the replicate and cell line. We decided to keep these proteins in the present build of the database to maintain maximum coverage. When replicates were merged for each cell line, the total numbers of proteins listed in the database for each cell line ranged from 5’709 (HCT-116) to 8’636 (HEK293). In an overwhelming majority of cases (97-98%) the accession code (AC) listed in the database was the first AC in the protein group in which the sequence was matched in the corresponding MaxQuant result table, showing that only few ambiguities due to protein inference in distinct samples are present. The low redundancy of the annotated UNIPROT sequence database used for identification plays a key role in this respect.

Between 59% and 84% of the proteins were detected as migrating as a single peak (Supplementary Table ST2). Also in a majority of cases the “highest peak” detected was clearly the major migrating species : the mean fraction of total signal accounted for by the highest peak ranged from 0.89 to 0.97 across all cell lines and replicates. While this value appears very high, it likely includes a bias, given that the numerous proteins that have a weak signal close to the limit of detection tend to be detected only once at a single position, resulting in a single peak. Still, when more than one peak was detected for a protein, the highest peak was, among all peaks detected, the closest to the expected MW (theoretical mass) for the protein in a majority of cases (61-72%). Overall, the data suggest that a majority of proteins migrate as a dominant single electrophoretic peak, which is often close to the expected mass. Although a majority of proteins present such an ideal behaviour, there is a significant number of proteins that deviate from it, either because they give rise to multiple peaks or because the main peak is at a MW significantly different from the theoretical MW calculated from the sequence. We address these cases in a later section.

Globally, distributions of highest peak MWs were very similar among individual cell lines and corresponded very well to the distribution of expected theoretical MW for the set of proteins identified. More importantly, the distribution was also very close to the global one calculated for the entire human proteome, suggesting that there was no major bias in our pipeline for or against proteins of a particular mass range (Supplementary Fig.S3A).

### Quality control : binning and maximum error in MW determination

In our workflow, all signals detected in a single gel slice are assigned the average MW of the centre of the slice. This results in binning of protein signals and thus an inherent technical error in MW determination. Locally, the maximum such error is thus related to the size of the MW range covered by each individual gel slice, which in turn is heavily dependent on the gel resolution at different MWs. We have determined that the bins corresponding to gel slices have a relatively constant size of 2-3 kDa in the range between 10 and 80 kDa but become larger at higher MWs, as can be expected from the gel separation pattern and from the fact that excised gel slices had a constant physical size (Supplementary Table ST3, Supplementary Figure S3B). The distribution of bin sizes also allows us to derive the maximum technical error due to gel cutting and binning, which is equal to the half of the slice size in kDa. In all samples, such error remained between 1% and 5%, depending on the MW. The best accuracy (smallest bins) was observed around 50kDa, which corresponds to the region of maximal protein density, while the lowest accuracy was observed at high MWs above 250 kDa (Supplementary Fig. S3B). To reduce this intrinsic error in MW determination we have on one hand tried to maximise the number of slices, while keeping a reasonable throughput. On the other hand, we hypothesized that the merging of data from replicate runs should reduce binning effects, because distinct gels being cut result in slices centred at slightly different MWs, defining different bins for each run. Histograms of highest peak MWs for the 3 replicates *vs*. the merged interpolated data illustrate the transition from a discrete distribution in the replicates to a more continuous distribution of MWs after merging, which resembles more closely the theoretical distribution calculated from the protein sequence (Supplementary Fig. S3C). In conclusion, while gel slicing results in inherent biases of MW determination, we think that such effects are significantly reduced by the combination of a sufficient resolution in gel slicing with the averaging of replicates.

### Quality control and global assessment of MW values (all proteins)

For a first evaluation of the global similarities of migration patterns measured in different cell lines, we calculated correlations of highest peak MW values across all replicates of all cell lines (Supplementary Figure S3D). Pearson’s R coefficients varied between 0.74 and 0.95, with nevertheless most values above 0.85. Interestingly, correlations intra-cell line were not drastically higher than between different cell lines. Averaged Pearson’s R coefficients for any particular 3 vs. 3 comparison (intra- or inter cell lines) were all greater than 0.80 (Supplementary Figure S3D). Next, to assess the precision of measurement of highest peak MW, we calculated coefficients of variation (CV’s) of this parameter. Across replicate measurements in the same cell line, the mean CV ranged between 5.3 and 6.4% (medians were between 1.8 and 4%). This suggests that MW measurements are reproducible for all cell lines analysed. The overall mean CV across the entire database, calculated using all replicates across all cell lines, was higher, at 10.1%. However when replicates were averaged first for each cell line and the averages used to calculate CV’s across cell lines, the final mean CV was of 7.1%, i.e. almost identical to the values intra-cell lines. This suggests that averaging within each cell line further improves precision. The fact that CVs for the highest peak remain moderate even between distinct cell lines is in line with the empirical expectation that, globally, the main species of most proteins (though by no means all) should migrate at very similar MW in multiple cell lines.

To further benchmark the overall correlation and precision of MW determination we used a set of 2688 proteins detected with a highest peak accounting for at least 70% of their total intensity in all replicates and cell lines (Supplementary Table S4). Analysis of these proteins that migrate predominantly as a single species and for which MW can be determined with high confidence yielded Pearson’s R correlation coefficients that were uniformly higher than 0.98 between all replicates and cell lines, indicating high correlations (Figure 3A). Once more, correlations between cell lines were almost as high as within replicates of the same line. Highest peak MW values between cell lines after interpolation and sum of the replicates were also very highly correlated, i.e. greater than 0.93. For this set of proteins, mean CV’s of MW values within individual cell lines were very low, ranging from 2.1% to 4.1%. When all replicates of all cell lines were considered for calculation, the CV was in the same range (3.2%)(Supplementary Table S4). The consistency of the measurements indicates that, for proteins detected reproducibly in more than one replicate, the value of MW can be mostly measured with high precision.

**Figure 3.**
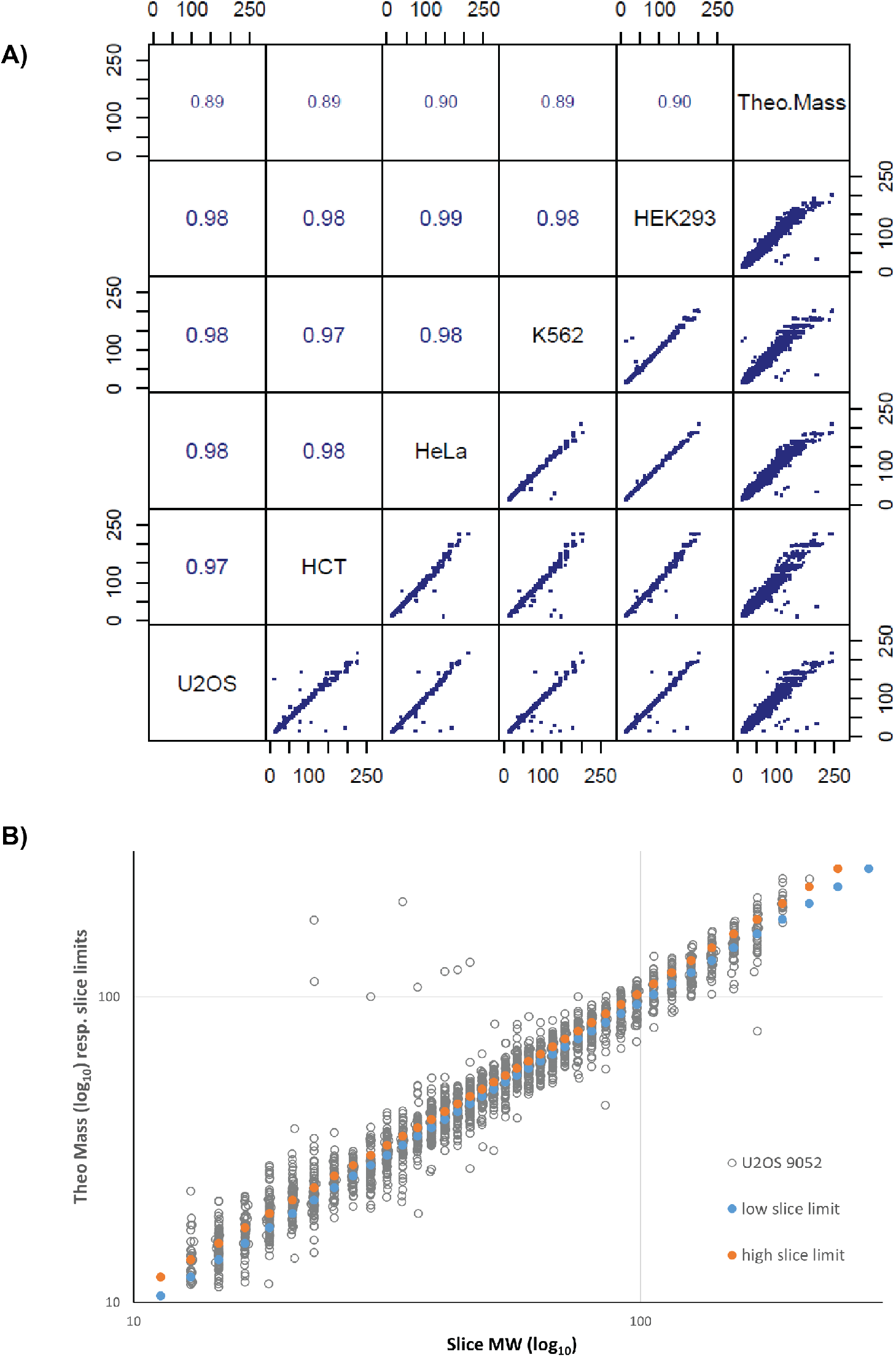
Correlation of MW measurements and evaluation of systematic errors. **A)** Correlation of highest peak MW values among cell lines and to theoretical MW for a set of 2’688 proteins detected in all cell lines as a single main band (Table S4). Each value results from interpolation and averaging of 3 replicates. Values in the upper portion of the plot are Kendall’s correlation coefficients for the corresponding comparison. Theoretical MWs (Theo.Mass) were calculated from the UNIPROT default sequence. Font size of correlation coefficient is scaled proportional to its value. Axis marks on scatter plots are values of MW in kDa. **B)** Evaluation of systematic errors due to binning in gel slices. In grey : plot of theoretical MW vs. gel slice center mass. Blue and orange : low and high slice limits vs. gel slice center mass. Data is for one representative dataset (U2OS, repl.1). MW scales are logarithmic (log10) but labels are in linear units.

### Deviation of measured MW from calculated MW

Interestingly, all correlations of experimentally determined MW values between cell lines and replicates were clearly higher than correlations of experimental MW to theoretical mass values (0.984 vs 0.968, Figure 3A). To determine if the deviation from the theoretical MW can be explained technically by the bias introduced by the slice-based binning we compared the fractional mass deviation (FMD) of proteins in individual replicates with the maximum error due to binning (Figure 3B, Supplementary Fig. S3B). We defined the FMD as the difference between the observed position of the main peak and the theoretical mass, divided by the theoretical mass. The plot obtained shows that binning bias due to gel slicing can account for only a small fraction of the differences observed between electrophoretically measured MW and the expected mass. The implication of all the observations above is that proteins can and do display highly reproducible migration, but also reproducibly deviate from their “ideal” migration behaviour predicted from the calculated molecular mass. This observation prompted us to systematically investigate possible intrinsic biochemical properties that could influence the electrophoretic migration of proteins (see below).

### Online database : interface and features

The database, that we codenamed PUMBA, is freely accessible online (https://pumba.dcsr.unil.ch/). The user interface has a welcome page with a search box that accepts UNIPROT identifiers or gene names and the possibility to choose the cell lines. If the protein of interest is matched, the default “lanes” view is western blot-like (Figure 4A) and shows for each cell line the merged pattern obtained from the three replicates. It is possible to show the data of the individual replicates with a click on the lanes. The user can adjust the grey level of the view to simulate the effects of a longer/shorter exposure on a physical WB (Figure 4A, right panel). This allows to see weaker bands that may correspond to modified forms or cleavage and degradation products.

**Figure 4.**
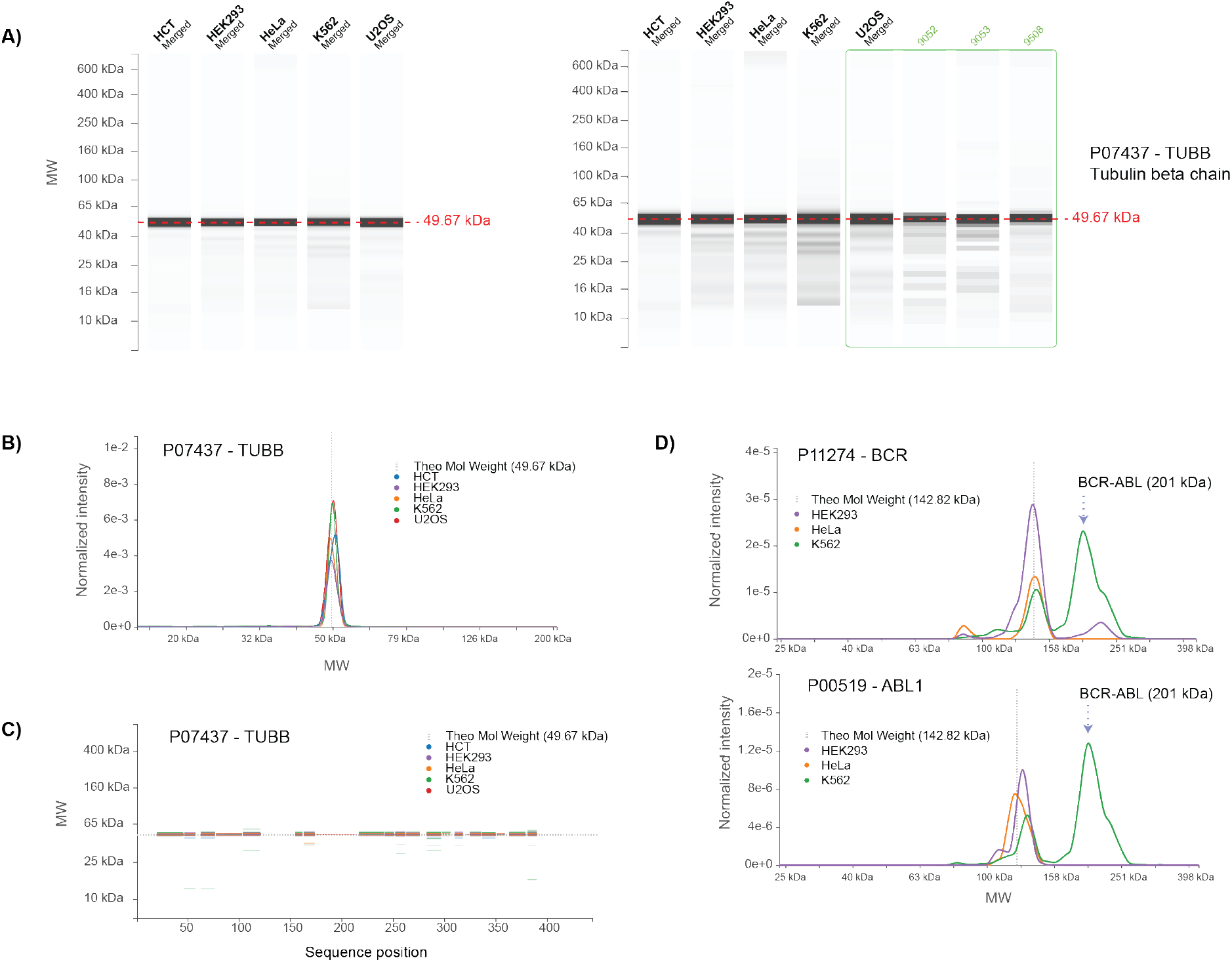
Graphical user interface of the PUMBA database. **A)** The default “Lanes” view is a virtual western blot. In this view it is possible to adjust contrast to visualize weaker bands as well as show data for individual replicates (right panel). **B)** The “Graph” view represent the same information but allows superimposition of traces for the individual lines, zooming and cursor-based MW display. **C)** In the “Peptides” view, peptide matches are shown in function of their location in the sequence and their identification at MW positions, also with the possibility of zooming and intensity-based thresholding. **D)** Graph views for proteins BCR and ABL1 are shown. The position of the BCR-ABL fusion protein expressed in K562 cells is indicated (arrow).

The second view is in a graph format (Figure 4B) and allows a more in depth comparison of the pattern among cell lines by superimposition of the traces. Here, too, it is possible to display the raw data for the replicates as pop-ups, to adjust the scale of the signal and to zoom in a region of interest (Supplementary Figure S4A). The intensity of each trace is based on the MS intensity as determined by MaxQuant, expressed for every protein as a fraction of the total intensity for all identified proteins. While not normalized by sequence length and probably not strictly quantitative, the intensity values permit a rough estimation of the overall abundance of a protein in a cell line. So for example, the peak for beta tubulin (TUBB) has a similar apex intensity in all 5 cell lines, between 2×10e-3 and 3.5×10e-3. By contrast, the tumour suppressor p53 (TP53), detected only in HCT-116 cells, has an apex intensity of 2.5×10e-6, i.e. approximately a thousand fold lower.

The third view (“peptides “) maps the identified peptides on the sequence of the protein on the horizontal axis as a function of the recovery of the same peptides in the gel MW range (y-axis) (Figure 4C). This unusual format of visualization offers the advantage of presenting the total information on the recovery of peptide fragments at all MWs. Like in the other views it is possible to filter the peptides shown by cell line but also adjust the display based on intensity, a feature that is particularly beneficial for interrogation of highly abundant proteins. Indeed, these are often detected in trace amounts at almost any MW, presumably due to degradation products and the gel migration trail, but only show high intensity signals at the position corresponding to the main band (Supplementary Figure S4B).

As an example of usage of the graphical interface we queried the database for the BCR-ABL protein. This chimeric polypeptide results from a chromosomal translocation (Philadelphia chromosome) and is a hallmark of K562 cells, isolated from a myeloid leukaemia patient. Three out of five cell lines show peaks corresponding to endogenous BCR (highest peak MW=144 kDa) and to endogenous ABL1 (121-131 kDa) (Figure 4D). But only in K562 cells the profile shows for both of these proteins an additional peak at 200 kDa that corresponds to BCR-ABL. Furthermore, the shape of the profile for the fusion protein in K562 cells is essentially identical, regardless of whether BCR or ABL1 peptides are being detected (Figure 4D, Supplementary Figure 4C). This example shows that the approach can detect different parts of the same sequence across the migration profile as well as species with unexpected migration behaviour.

### Molecular properties influencing gel migration

After establishment of the database and quality control of the content, we mined the data to try and identify protein properties that could influence electrophoretic migration. For this we considered the above described set of 2688 proteins that are ubiquitously detected mainly as a single dominant (>70% of intensity) peak, together with their FMD values. Since this analysis only considers the main peak, it will highlight only deviations from the ideal migration that affect the bulk of the pool of a protein. One thus expects to detect only effects due to either intrinsic properties of the sequence or to constitutive, high-stoichiometry post-translational modifications.

Distributions of average FMD values in this set of proteins were very similar among cell lines and were centred close to zero with interquartile ranges of 0.12 and a majority of values between −0.2 and +0.2 (Supplementary Figure S5A). We first examined the correlation of FMD with isoelectric point (pI) and hydrophobicity. A plot of FMD vs. pI does not suggests a clear, global correlation between protein net charge and slower or faster migration (Figure 5A). Only a few proteins with extreme pI values show a tendency to migrate abnormally (mostly slower, i.e. positive FMD values).

**Figure 5.**
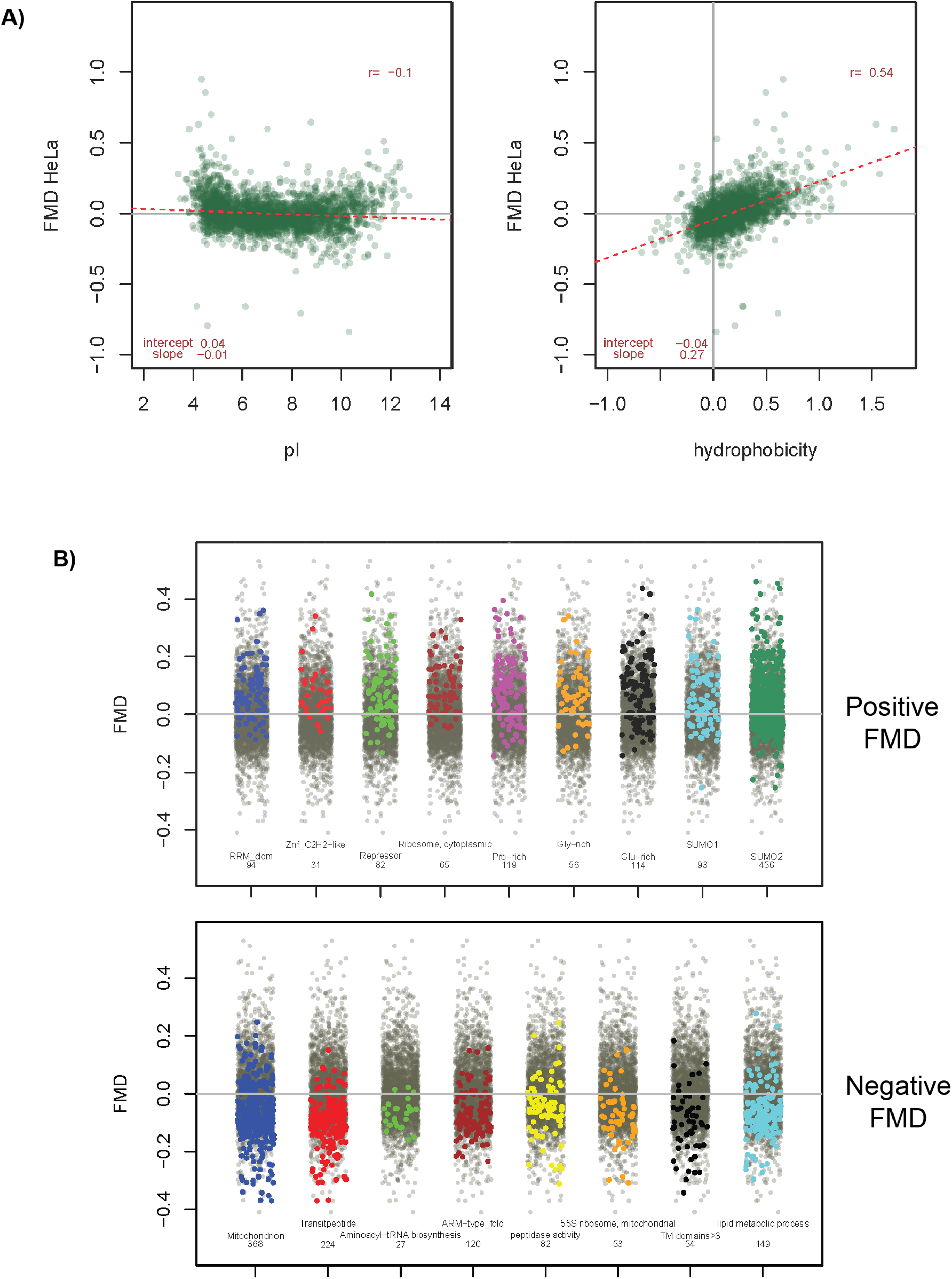
Correlation of Fractional Molecular Weight Deviation (FMD) with molecular properties and functional annotation. A set of 2688 proteins detected in all 5 cell lines with a main single peak accounting for more than 70% of total signal was used for the analysis. Data shown is for HeLa cells, see Supplementary Fig.S5 for all other lines. **A)** FMD values as a function of pI (left panel) and hydrophobicity (right panel, calculated by the method of Hopp-Woods. **B)** stripchart plots of selected annotation terms enriched for positive (upper panel) or negative (lower panel) FMD values in at least four out of five cell lines. Values for proteins with the specific term are highlighted in colour against the total set in gray. Upper panel terms are : RNA recognition domain (Interpro); Zinc finger-CH2 type domain (Interpro); Repressor (UNIPROT keyword); Ribosome, cytoplasmic (Corum); Pro-rich,Glu-rich,Gly-rich (compositional bias, UNIPROT); SUMO1/2 (UNIPROT PTM). Lower panel : mitochondrion (UNIPROT keyword); transit peptide (UNIPROT keyword); Aminoacyl-tRNA biosynthesis (KEGG); ARM-type fold (Interpro); peptidase activity (GOMF); 55S ribosome, mitochondrial (Corum); TM domains>3 (UNIPROT annotation); lipid metabolic process (GOBP). The position of points for the background total set (grey) are randomized by the plotting algorithm and thus appear different in each plot.

On the other hand, a correlation was observed between FMD and hydrophobicity (Figure 5B), suggesting that very hydrophobic proteins migrate on average faster than expected (FMD<0) and show MW values lower than predicted based on their sequence.

We then carried out 1D annotation enrichment analysis on the considered set of 2688 well-detected proteins sorted by FMD values. The categories considered included pathway and Gene Ontology terms (KEGG, GOBP, GOCC, GOMF) but also terms related to structural and sequence features from several sources (Interpro, Corum) as well as terms derived from manual UNIPROT/Swissprot curation (e.g. “keywords”). Even considering only terms statistically significant in at least 4 out of 5 cell lines, a number of annotation terms emerged that correlated with positive or negative FMD values (Supplementary Figure S5B). Results for global functional annotation terms (GOBP, KEGG) suggested that ribosomal and spliceosomal proteins and in general proteins involved in RNA processing and transcriptional regulation have positive FMD values, i.e. migrate slower than expected. On the other hand, proteins involved in several mitochondrial metabolic pathways and oxidative phosphorylation seemed to have negative FMD (Figure S5B,C), i.e. migrate faster than expected. Similar trends emerged from annotation focusing on localisation (GOCC, Corum).

These broad categories encompass very heterogeneous groups of proteins with mixed molecular properties. In the subsequent analysis we assumed that deviation of migration from an ideal behaviour under denaturing conditions must be due to either properties of the primary structure (sequence) or else to PTMs that impact mass, charge or hydrophobicity independently of 3D structure. We thus focused on more structural annotation to identify features that clearly correlate with positive or negative FMD and whose categories show minimal overlap. We then verified if the positive/negative FMD averages observed for broader functional or topological groups were due to presence of a subset of these more structurally-based FMD “outliers”. For example, mitochondrial proteins, as defined by GOCC or UNIPROT keywords, have on average negative FMD values (Figure 5B). Based on our initial assumptions, we speculated that this is caused by the proteolytic removal of transit peptides that occurs during chain translocation through the mitochondrial membrane. Indeed, a set of 368 proteins with the keyword “Mitochondrion” has an average FMD of −0.08 (for HeLa cells), but when 223 proteins in this group annotated as having a transit peptide are subtracted, the remaining 145 protein set has an average FMD of −0.02, very close to the global average. This suggests that for a majority of integral mitochondrial proteins the observed lower-than-expected migration is, unsurprisingly, due to transit peptide removal. This is indirectly supported by the comparison of cytosolic ribosomal proteins, which as a group have a strikingly positive FMD, with mitochondrial ribosomal proteins, which have negative values after being synthesized in the cytosol and imported in mitochondria (Figure 5B). Using this approach, we found that positive FMD was associated with proteins having Interpro RNA Recognition Domains (RRM_dom), a set that overlaps in part with that of proteins annotated with a “nucleotide-binding alpha/beta plait”. A similar but only minimally overlapping group was characterized by presence of classical Zn-finger domains (Interpro ZnF C2H2). Further on, proteins with a (transcriptional) “repressor” activity (UNIPROT Keyword) were also positive, but only minimally overlapped with the groups previously described. In turn, “cytosolic ribosomal proteins” (Corum) stand out as very hydrophilic, high-migrating group but did not contain a significant amount of the annotations listed above.

Surprisingly, amino acid compositional bias (from UNIPROT keyword annotation) appeared to be always correlated with positive FMD values, independently of the amino acid that is overrepresented (Supplementary Figure S5, Fig. 5B). Major groups with significant shifts were Pro-rich but also Glu- and Gly-rich proteins.

We specifically asked if modification by cross-linking to ubiquitin family members, which can induce large MW increases, resulted in a tendency to positive FMD. Surprisingly, only a mild correlation was observed with sumoylation by SUMO-1 and SUMO-2. This could be due to several factors, including the incompleteness of the annotation but also the fact that such modifications (like many other PTMs) are often sub-stoichiometric and would not be covered by this analysis that only takes into account a shift of the bulk pool of each protein.

Concerning negative FMD shifts, many functional categories (TCA cycle, cellular respiration, cellular ketone metabolic process, .., Supplementary Figure S5) can be explained by the mitochondrial localization of the corresponding proteins. Other distinct, non-overlapping groups with negative FMD included the majority of aminoacyl-tRNA synthetases as well as a quite large group of proteins with an ARM-type fold domain (Interpro annotation). Unsurprisingly, proteins with a peptidase activity (GOMF) were among the ones with the strongest negative FMDs, likely due to frequent (auto-)proteolytic cleavage that is essential to production of the active form (e.g. TPP1 or Calpain catalytic subunits CAPN1/2). Proteins with several (>3) transmembrane domains also appeared to migrate clearly below their expected mass (Figure 5C), in line with their marked hydrophobicity (average −0.19 for HeLa cells). Excluding (auto)proteolytic events, only few PTMs appeared to correlate with significant (positive or negative) FMD values and these were mostly small groups (Supplementary. Figure S5B), e.g. ADP-ribosylserine (10 proteins, positive) or N6-(pyridoxal phosphate)-lysine (14 proteins, negative FMD).

In conclusion, we could identify numerous groups of proteins, defined by annotation terms, which show systematic shifts in electrophoretic mobility relative to their theoretical mass. While it is possible to identify some trends, it remains difficult to pinpoint conclusively the factors causing such shifts because proteins have with many and multiform domains, features and properties. The existence of unknown PTMs can also not be ruled out. Furthermore, biological annotation is redundant, possibly incomplete, and contains many overlapping terms.

### Individual proteins with noncanonical migration patterns

After global exploration of properties that possibly affect electrophoretic migration, we tried to assess how well the database can capture the behaviour of individual proteins with more complex patterns and/or proteins with MW that strongly deviates from the predicted value reported in the database. We looked for known cases representative of three types of events : post-translational cleavage, differential splicing and conjugation.

### Proteins with unusual migration patterns due to cleavage : PTPRF

Some human proteins undergo constitutive cleavage to generate the mature form. Such proteins may or may not present a remnant peak/band at the expected full length (FL) MW, and should show signals at the MWs of the known fragments. In addition, the peptide sequence coverage at the different positions in the gel should be consistent with known position(s) of the cleavage site(s). A representative example is the receptor-type tyrosine-protein phosphatase F (PTPRF), a 1907 amino acid-long type-I membrane protein with a single transmembrane domain (TM). The protein is cleaved extracellularly adjacent to the TM domain during export^27^. In the PUMBA database, PTPRF is well detected in both HeLa and HEK cells as the N- and C-terminal fragments, respectively at 132 and 78 kDa (Figure 6A). The peptide map in PUMBA locates the cleavage site between pos. 1150 and 1180 (numbering based on full sequence). This fits with previous reports^27^ locating the cleavage site in the RRRRR motif located at position 1174-78. By inspection of the database we could verify detection of known cleavage events for several other proteins (P52948, P07686, P07339, P10619, P07858, Q9H7Z7, P46821). However, a comprehensive search for chain cleavages would require an in-depth study.

**Figure 6.**
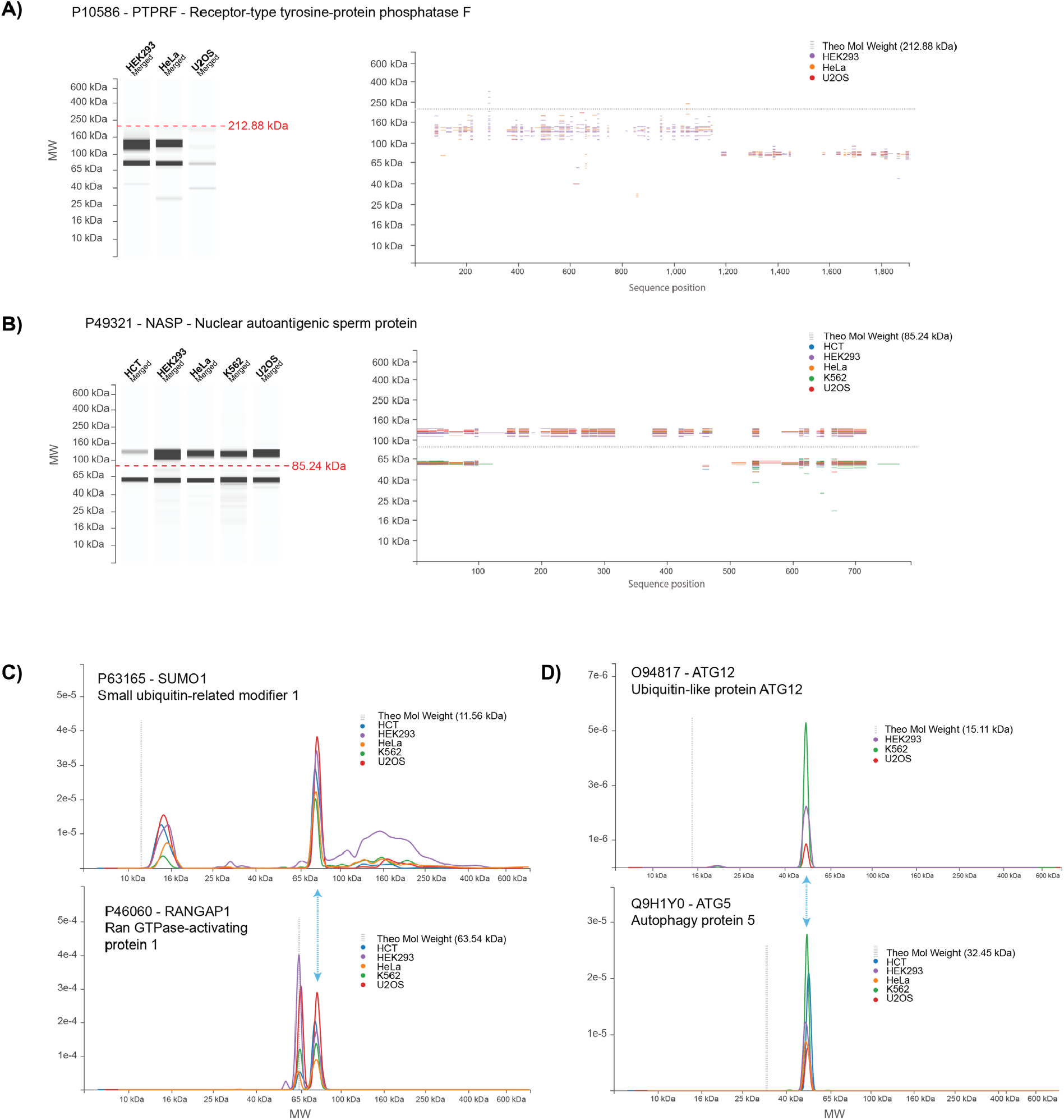
Database entries for representative proteins with MW peaks strongly deviating from their theoretical value. Examples due to constitutive cleavage **(A)**, differential splicing **(B)** and conjugation **(C**,**D)** are described in the text.

### Proteins with unusual migration patterns due to differential splicing : NASP

Expression of a different splice variant can cause important changes in protein MW. In this respect it has to be noted that, unless clear information is available, the UNIPROT database reports by default the longest splice variant as the default protein isoform. As an example we focused on NASP (Nuclear autoantigenic sperm protein P49321 (NASP_HUMAN), a widely expressed histone-binding protein with chaperone properties. The full length canonical isoform of NASP has an expected MW of 85 kDa and 4 coiled-coil domains of 152 AA in total, which constitutes almost 20% of its sequence. Moreover a large portion of its sequence is rather acidic (Glu-rich), a fact linked to its histone-binding properties. We detected human NASP in all cell lines as two bands, migrating respectively at 120-125 and 60 kDa (Figure 6B). Peptide map coverage of the 120 kDa band is extensive, suggesting presence of the full sequence. Based on our previous findings, we speculate that the coiled-coil domains and Glu-rich sequence could account for the “excess” apparent MW of 35 kDa relative to the expected MW. An equally strong band is found at 60 kDa, which displays a markedly different peptide map, with a gap between AA 123 and 489. This sequence coverage fits very well to NASP isoform 2 (P49321-2), a splice variant missing residues 138-476 and has an expected MW of 48 kDa.

Isoform 2 retains only 2 of the coiled-coil domains present in the full length sequence as well as much less of its acidic stretch, which could correlate with its reduced “excess” MW of only 12 kDa. Finally, the intensity data in PUMBA allows a rough comparison of the relative ratio of the two (60 and 120 kDa) isoforms, which appears to vary significantly among cell lines. The two species detected correspond to the previously described isoforms sNASP and tNASP (somatic and testicular NASP) that have been characterized in mouse tissues. Both variants are known to be widely expressed, and our data confirm in human cell lines the previous findings in mouse^28–30^.

### Proteins with unusual migration patterns due to conjugation to other proteins: SUMO-1 and others

Conjugation to proteins of the Ubiquitin family is a major regulatory mechanism resulting in generation of species with altered MW. SUMOylation (by SUMO-1) notoriously regulates a number of cellular processes^31^. The migration profile for SUMO-1 in our database shows a peak at 15 kDa that probably corresponds to the free monomer, which has an expected MW of 11.56 kDa (Figure 6C). The main peak for SUMO-1, however is found around 77 kDa. Although SUMO-1-ylation affects a number of proteins, it is known that the Ran GTPase activator RANGAP1 is a major SUMOylation target in the cell^32^. In PUMBA, the main RANGAP1 peak is found at 63.5 kDa, in very good agreement with the theoretical mass of the protein (63.54). But a peak of similar intensity is found at 76-78 kDa, which overlaps perfectly with the intense SUMO-1 peak (Figure 6C). Furthermore, the hierarchy of intensities among cell lines for this peak is highly similar in the RANGAP1 and SUMO1 profiles with the HCT-116, U2OS and HEK293 cells on top and HeLa and K562 cells having lower values. This, together with the high similarity of the profiles, suggests that it is the same species, albeit identified and quantified on different peptides. Besides the monomer and the RANGAP1 peak, the SUMO-1 profile shows a weaker, diffuse signal on a large range of molecular weights as expected for the ensemble of SUMOylated proteins in the cell. A similar but more diffuse profile can be observed for Ubiquitin (e.g. P62979 in PUMBA, see https://pumba.dcsr.unil.ch/), for which it is however difficult to infer unambiguously a dominant target. Highly specific conjugation was on the other hand clearly seen for the ATG5-ATG12 pair (Figure 6D). Overall, our approach offers a way to survey the MW distribution and intensity of Ub family-conjugated species that should be less biased than antibody-based detections. At least for ubiquitin, the latter show limitations due to the different recognition of diverse types of linkages by the various antibodies available^33^.

The three examples above show that, if a sufficient sequence coverage is available, data in our database can accurately recapitulate known complex migration patterns and map processing or splicing events when these affect significant portions of the sequence.

### Possible novel processing or differential splicing events and database annotation

Next we hypothesized that the PUMBA database can provide hints for as yet un- or poorly characterized processing or splicing events. In practice, the data is especially amenable to identify either cleavage events or splice variants that are shorter than the canonical one. This is a rather common case, since the canonical sequence listed in UNIPROT is almost always the longest isoform. In both cases, evidence should be based on a lower-than-expected highest peak MW in combination with a partial peptide sequence coverage that fits well the observed molecular weight. We thus carried out manually a (non-comprehensive) survey of the database, examining more in detail proteins with absolute values of FMD greater than 0.2. A certain number of candidates emerged, which present patterns compatible with differential splicing or processing distinct from the information presented in UNIPROT (Table 1). While the evidence in the PUMBA database alone cannot be taken as conclusive proof, we believe it may constitute a starting point toward discovery and elucidation of processing events or differential splicing.

**Table 1.**
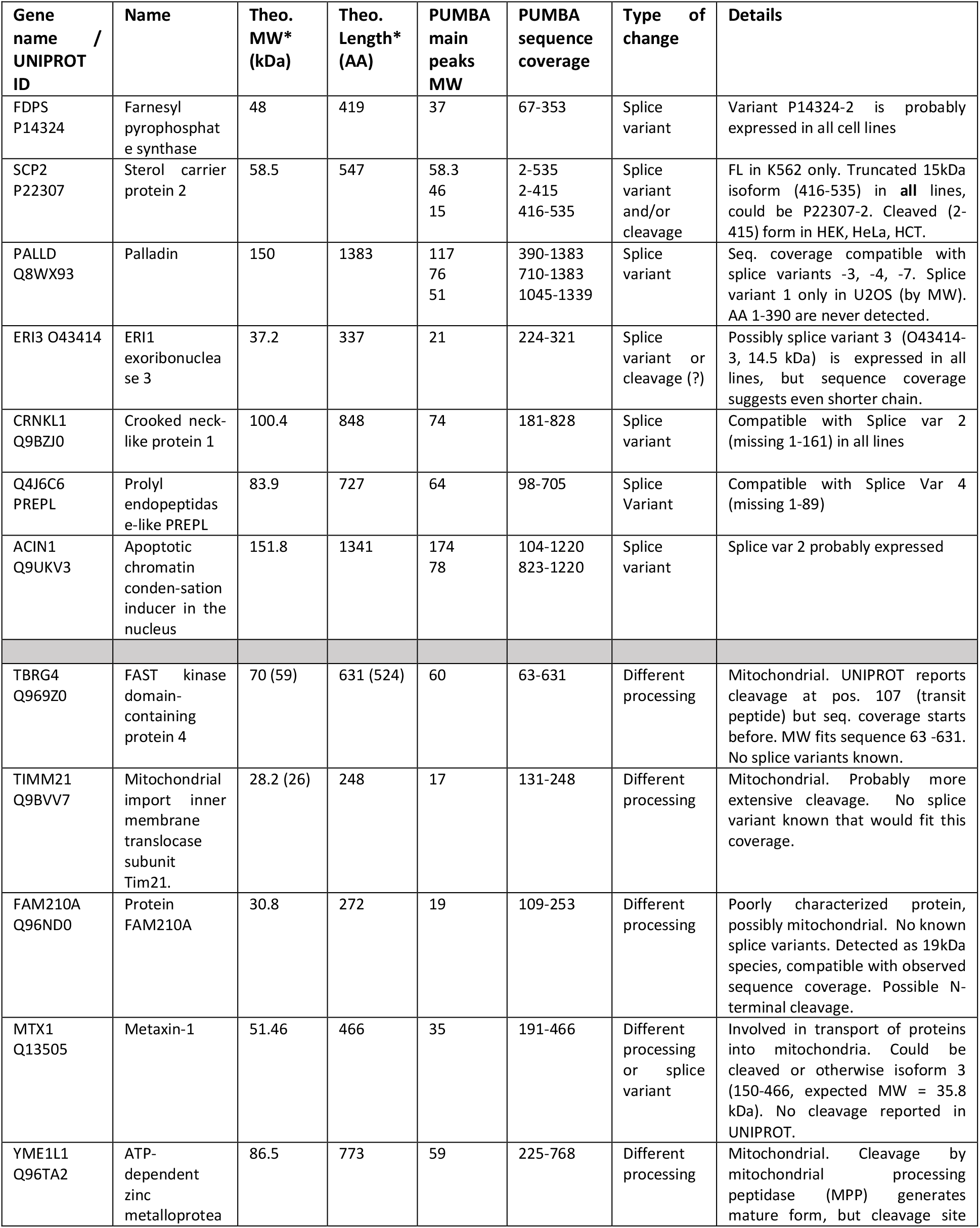
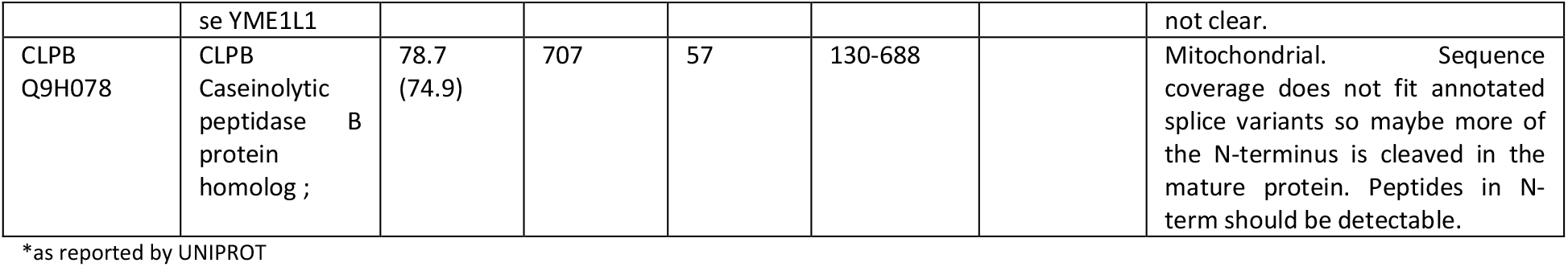
Proteins with MW differing from sequence-predicted values : new or poorly characterized cases.

## Discussion

We present here a first version of a repertoire of reference electrophoretic patterns for human proteins in a set of commonly used cell lines. The usage of a geLC-MS approach to build migration maps is not new, but most previous efforts aimed at studying specific biological phenomena that impact protein migration by comparing two or more conditions^11,12,14,34,35^. In a different context, Edfors et al ^36^ showed that the approach can be used for antibody validation, using the human cell lines RT4 and U-251 that were also used for large scale Western Blot in the Human Protein Atlas project. Only Yang et al ^15^ constructed a freely accessible, online “virtual Western” database of protein migration for the mouse collecting duct cell line mpkCCD and used it to explore migration behaviour as a function of properties. Our work expands on these previous efforts but with a different scope, as we aimed to create a reference resource covering human cell lines widely used in research labs. To do this, we needed a framework to i) generate accurate, reproducible MW values and migration patterns and ii) make a maximum of information easily accessible. The resource should also be expandable to accommodate future additional datasets.

Our approach using internal calibration of migration yields reproducible and accurate electrophoretic MWs for thousands of proteins. To our knowledge, this is the first time that gel electrophoresis patterns for a large portion of a proteome are determined and can be compared across replicates that were run months apart as well as among different cell lines. In fact our data show that, if the experimental setup is constant, protein MWs can be determined accurately and reproducibly with very low CV’s. We believe that this is only possible thanks to the large amount of information generated by MS, together with the use of internal global recalibration rather than relying on external migration standards. The calibration curves obtained for our gels show that, globally, most of the proteome behaves “as expected”. At the same time, the MW of most proteins deviates slightly from the expected value, with in some cases much larger discrepancies. Relying on a set of 5-10 individual proteins for obtaining accurate MWs would be intrinsically flawed, as every one of these proteins would migrate with a distinctive deviation from the expected value, creating a sparse and somewhat distorted MW calibration curve. In contrast, fitting an average pattern based on several thousand different polypeptides has a much higher chance to yield a robust and accurate calibration curve since individual protein properties are averaged out. The calibration curve obtained, thanks to the density of its points, can compensate much more effectively for local migration differences intrinsic to every gel run and allows a much better alignment of different runs.

We showed that a majority of the proteins that are present as one dominant gel band display a MW that is highly similar across a range of different human cell lines. This suggests that protein migration is intrinsically reproducible, at least if the reagents and electrophoretic conditions are kept reasonably constant. While we did rely on commercial pre-cast gels and buffers, we made no special efforts to use reagents originating from the same batch. As mentioned previously, we successfully aligned gel runs that were carried out months or years apart, using different batches of gels.

At the same time, while a majority of proteins migrate close to their theoretical MW calculated from the sequence, small to medium deviations from such ideal behaviour are pervasive. Our analysis of correlation of physical properties and annotation terms with the FMD values revealed some trends, e.g. faster migration of hydrophobic proteins or slower migration of proteins with large regions with compositional bias. Our finding are in very good agreement and expand those of Yang et al.^15^, for example on the effects of hydrophobicity and cleavage on migration. Although the physicochemical phenomena underlying these effects are not always clear, knowledge of these biases can help rationalize migration behaviours of proteins not covered in this survey and possibly also in other species.

Our preliminary exploration of individual proteins with MW drastically different from the expected value highlight proteolytic processing as one of the main causes of unexpected (faster) migration, especially in proteins located in mitochondria or other subcellular compartments. It also stresses the fact that the details of constitutive proteolysis linked to chain maturation are still only partially characterized for a number of human proteins. Similarly, our data suggests that in a number of cases the dominantly expressed splice variant is likely not the longest isoform typically listed as the default one by UNIPROT (Table 1). While our findings by themselves cannot be fully conclusive, we often observe almost full agreement across the 5 cell lines, supporting the idea that such variant expression patterns are widespread and biologically relevant. Determining which splice variant is preferentially expressed can be very challenging for genes with complex exon/intron structure. Many variants differ by only a few codons/amino acids, which can be or not covered by mRNA or peptide sequences. Also, formally validating a shorter variant requires proof of absence of part of the sequence (by definition a tricky question). The conservative choice made by the UNIPROT curators to list the longest isoform as the default one is understandable, yet the question of splice variant expression is a crucial one and will have to be addressed soon^37^. Transcriptomics data will presumably play a major role in this task^38^, but the pervasiveness of translation regulation and alternative start sites in eukaryotic systems implies that protein information will be essential for validation. We argue that accurate and reproducible MW values, together with peptide coverage and the knowledge of sequence properties that impact migration, can be important criteria that, combined with mRNA sequences, can lead to reliable assignment of splice variant presence/absence.

The quality and completeness of information in the PUMBA database is limited by technical factors well known in MS-based proteomics. First, for proteins expressed at low level, data is sparse and sequence coverage is limited. Aiming to expand the database, the most immediate solution to this problem will be to employ newer MS instrumentation, which will increase both depth and peptide coverage. A different challenge is posed by sequence regions that are difficult to map because they contain either too many or too few trypsin cleavage sites, resulting in peptides that are either too small or too large for detection and thus constitute false negative hits. We plan to enhance the sequence display in the “peptides” view to flag such problematic stretches, as an aid to interpretation of the data. A possible improvement for sequences with low K/R content could be to complement the data with second pass, semi-tryptic searches, which in our experience can often reveal partial sequences due to nonspecific trypsin cleavage or peptide in-source decay. Such searches should be performed with special parameters, such as tight mass tolerances and considering only matches to proteins already identified with full trypsin, to avoid false positive hits.

In the near future, we plan to expand the database further by adding more human cell lines corresponding to other tissue types. We also plan to expand significantly the mouse section of the database (not discussed here), which at the moment contains only data for NIH 3T3 fibroblasts, by analysing several tissue extracts of relevance for the research community. Analysing entire, extensively fractionated gel lanes is time-consuming. Here too, the increased measurement speed of new MS instruments will help us to accelerate the rate of data acquisition by employing shorter LC-MS gradients.

### Impact

Ultimately, we believe that the information in our database will have an impact at two levels. First, it will be useful for investigators that use SDS-PAGE and WB on a daily base to work on human proteins. Accurate and reliable MW data, together with peptide maps, will allow them to critically examine the information provided by Ab suppliers in data sheets of western blotting antibodies. In these documents, the MW of the band attributed to the target protein can often be derived only inaccurately. This in turn generates uncertainties when comparing one’s own WB with the data sheet, whenever more than one band is detected or the signal is weak. Moreover, if the protein of interest is found in our database, investigators will be able to use one or more of the easily available cell lines we analysed as positive (or negative in case of proven absence) controls. The high level of consistency of migration that we could observe for most proteins across cell lines suggests that, at least in normally growing cells, MW in these lines can be determined reproducibly. For molecules that undergo gel shift-inducing modifications (e.g. cleavage), it will be possible to try to explain and rationalize diverging patterns obtained with antibodies raised against different epitopes, based on the peptide maps in PUMBA. When, as often the case, the information on the antigen used for Ab generation is missing, the peptide pattern measured combined with MW can often help to formulate hypotheses on possible epitopes to try and explain the observed band patterns.

Second, we think that the availability of our database could contribute to the global mapping of human proteoforms. The human proteome is believed to be far more complex than the size of the genome can suggest, due to differential splicing and post-translational modification events which expand the number of actual molecular species. Mapping the human proteoform landscape, even only partially, will be a major research task for the next decade^37^. Some recent efforts have been aimed at developing tools for the unbiased identification of distinct proteoforms^39,40^ based on peptide intensities from large scale MS data. While these tools are promising, connectivity information is often lost in bottom-up proteomics. We argue that integration of reliable molecular weight and migration data from several distinct biological samples will be very helpful for defining and validating existence of structural proteoforms. This will require novel software tools to analyse the content of our database, as well as extensive correlation to other types of information already available in protein knowledgebases such as UNIPROT. Finally, we estimate that the greatest benefit of this database will come from its use by the community at large as a protein resource that effectively completes the existing ones.

## Acknowledgements

Many thanks to Severine Lorrain for critical manuscript reading.

## Funding

This work was funded by the University of Lausanne.

## Notes

https://pumba.dcsr.unil.ch/

### Competing Interest Statement

The authors have declared no competing interest.

