## Supplementary Figures S1-S5 for "A database of accurate electrophoretic migration patterns for human proteins in cell lines"

### Figure S1

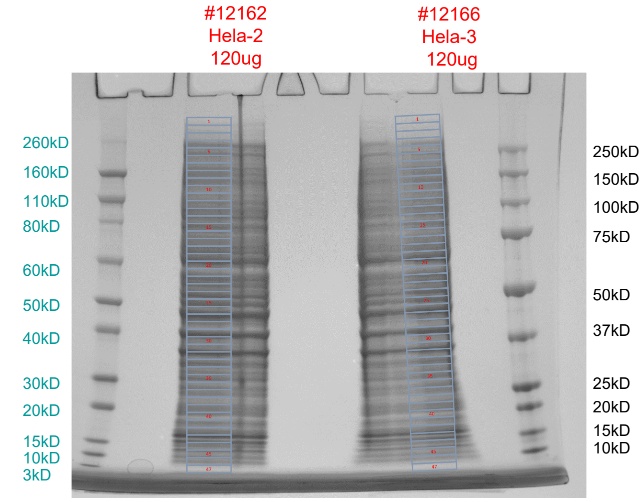


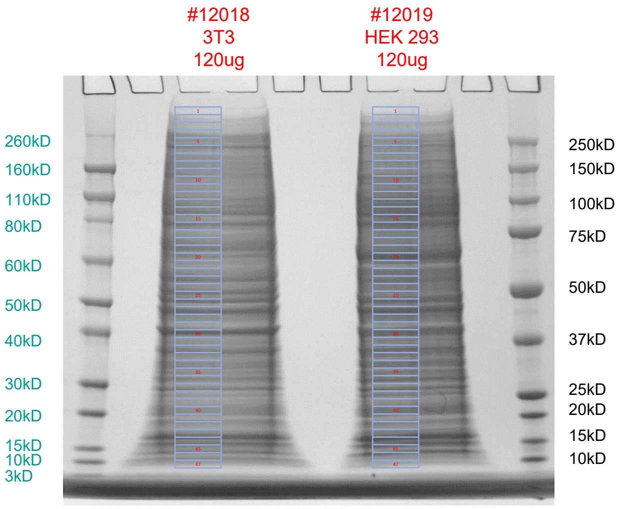


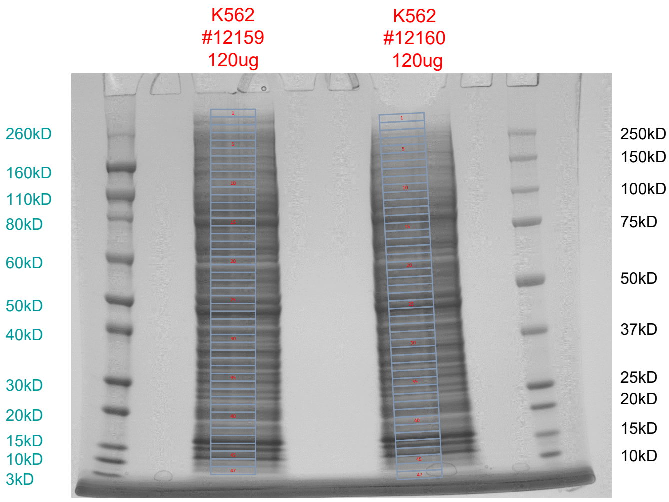


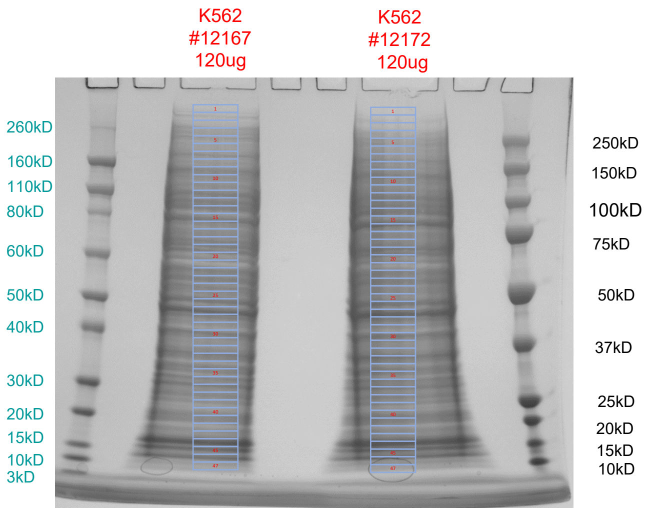


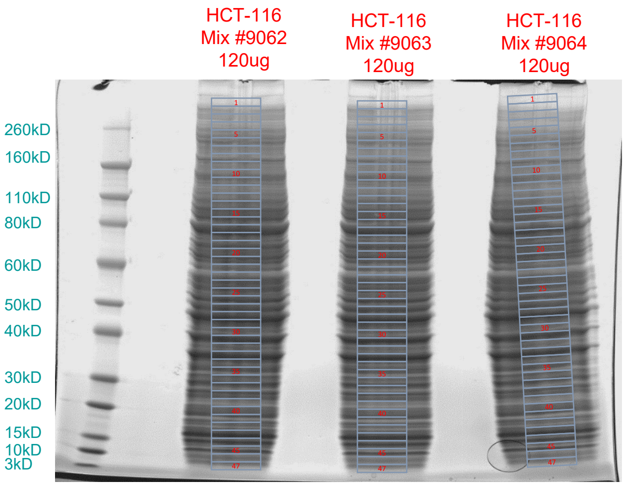


Representative gel migrations for the 5 human cell lines as well as mouse 3T3 cells. All cells were harvested by scraping in FASP buffer (4% SDS, 100mM DTT, 50mM Tris pH 7.5). Lysates were heated at 95C for 5 min, sonicated 30s and migrated on Invitrogen NuPAGE™ 4-12% Bis-Tris gels using MOPS migration buffer. Staining was by Coomassie Brilliant Blue G250.

### Figure S2

**A)**

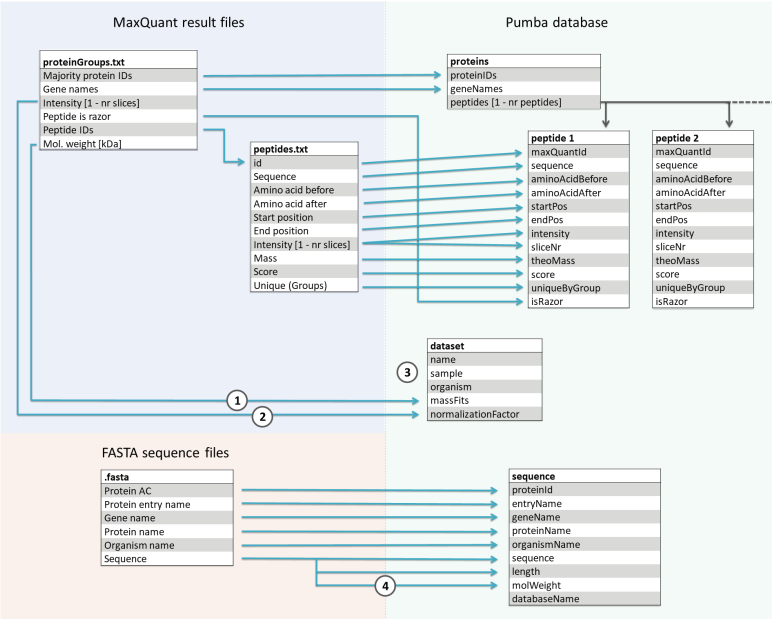


Import of fields from MaxQuant tables to construct the PUMBA database. Every "majority ID" is taken and compiled in the database together with its identification and quantitation parameters. For individual steps 1-4 in the scheme :

(1) A normalization factor is calculated by summing up all protein intensities from the dataset considered. This normalization factor is later used to compute the total protein intensities from the individual peptide intensities.

(2) The molecular mass of each slice is estimated by fitting a third degree polynomial function to a plot of theoretical protein masses against its slice position in the gel.

(3) The name, sample source and organism of a dataset have to be set upon sample upload.

(4)The theoretical molecular weight is calculated from the amino acid sequence

**B)**

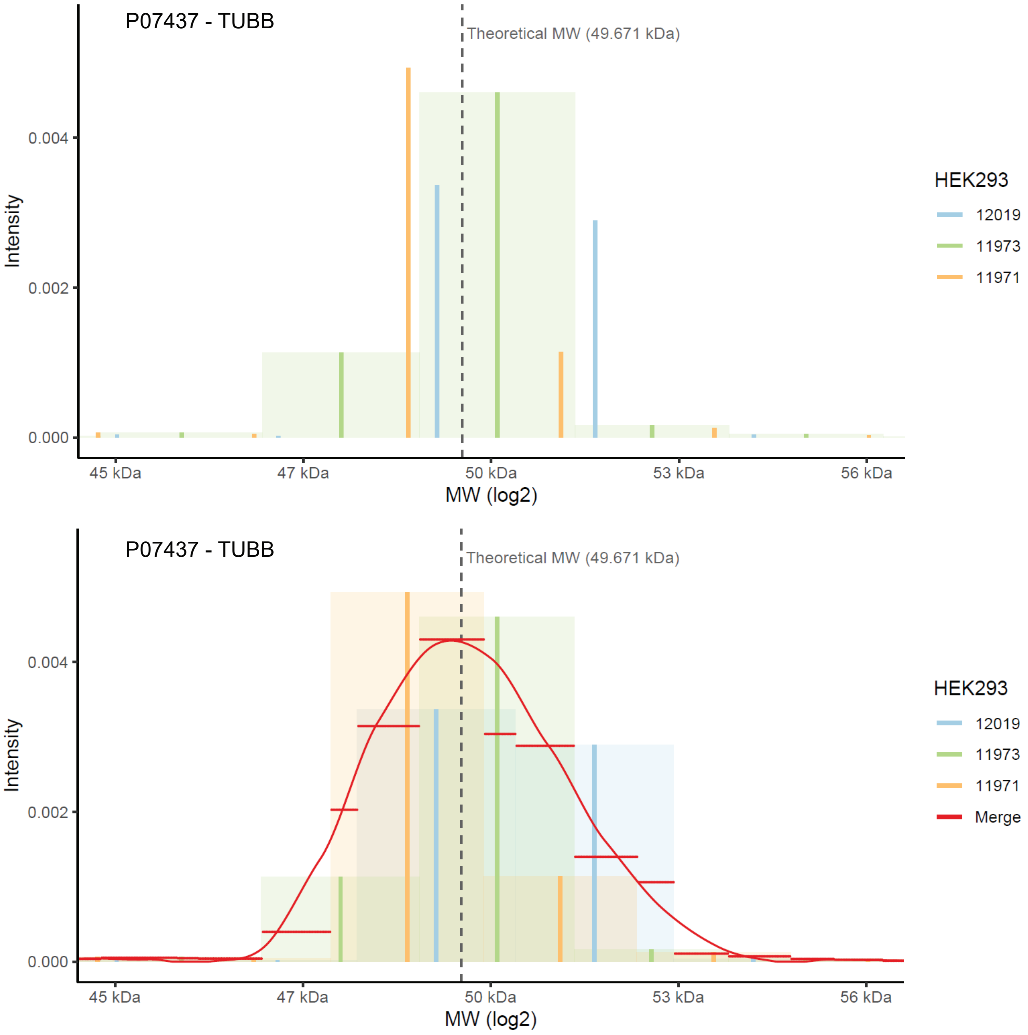


Computational strategy for interpolation, alignment and averaging of replicates. The raw, discontinuous signal for the protein TUBB is shown for three replicates of the same cell line. Each vertical bar corresponds to the intensity at the centre of one gel slice. A box is created which has the width of the gel slice and the intensity of the total intensity in the slice. This box is then split in 100 microslices along the MW coordinate (upper panel, shown only for the green replicate). The MW of each microslice is calculated using the fitted polynomial function. Microslices are then matched across replicates (by the closest MW). The intensity of matched slices is averaged, creating a step function (horizontal red lines, lower panel). This is then smoothed by a Loess algorithm to give the final, composite profile for the protein of interest. For representing individual replicates, as in the "Graph" interface, the box functions created can also be directly smoothed by Loess and plotted.

### Figure S3

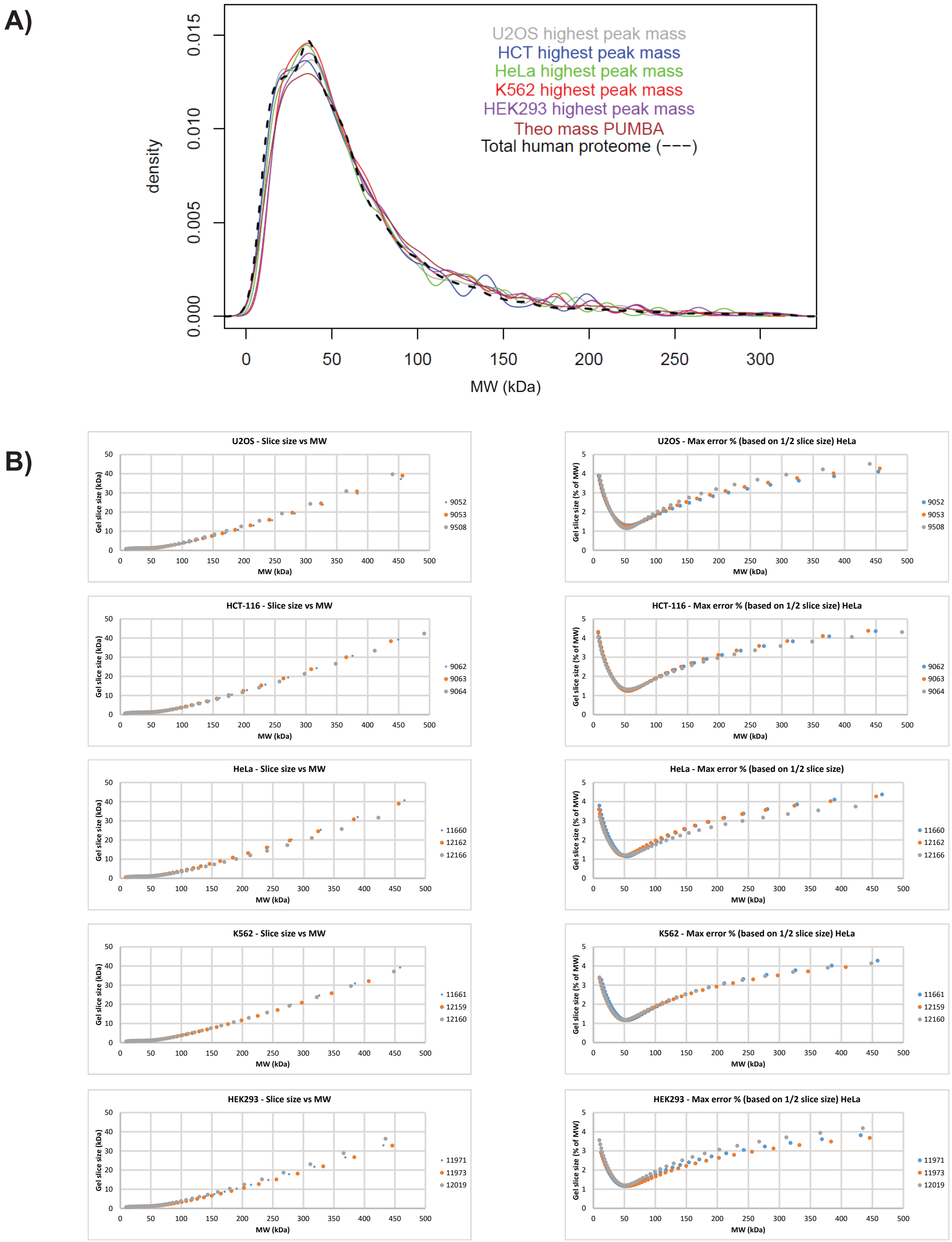


Global quality control : MW distribution and evaluation of systematic errors due to gel cutting **A)** Distribution of measured molecular weights in the 5 cell lines analysed, of their theoretical values and the total human proteome. Values of highest peak for all cell lines after averaging the 3 replicates are represented (Table S2, 10’187 proteins), together with their corresponding MWs calculated from the sequence (brown) and the distribution for the entire set of human annotated proteins from UNIPROT (20’371 sequences)(dashed line). Default parameters of the density function in R were used for the plot. **B)** Left column: absolute slice size as a function of MW. Right column: maximum errors due to slicing, equivalent to half slice size, shown as percentage of MW. Numbers correspond to labels of individual replicates (See Supplementary Table ST1).

Figure S3 (continued)

**C)**

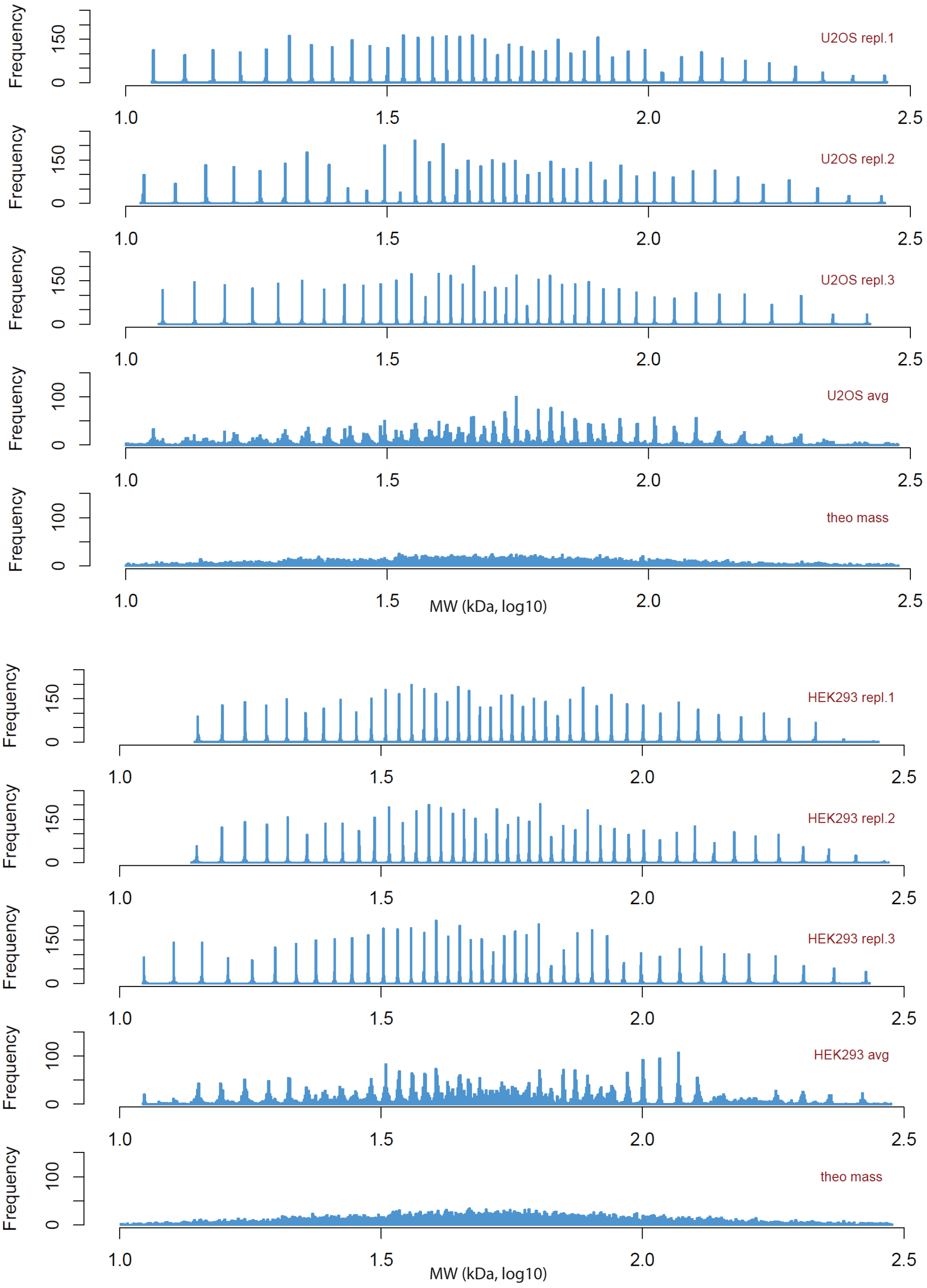


**D)**

| U2OS.9052.highest.peak.mass | U2OS.9053.highest.peak.mass | U2OS.9508.highest.peak.mass | HCT.9062.highest.peak.mass | HCT.9063.highest.peak.mass | HCT.9064.highest.peak.mass | HeLa.11660.highest.peak.mass | HeLa.12162.highest.peak.mass | HeLa.12166.highest.peak.mass | K562.11661.highest.peak.mass | K562.12159.highest.peak.mass | K562.12160.highest.peak.mass | HEK293.11971.highest.peak.mass | HEK293.11973.highest.peak.mass | HEK293.12019.highest.peak.mass | T: Name |
|---|---|---|---|---|---|---|---|---|---|---|---|---|---|---|---|
| NaN | 0.945 | 0.904 | 0.911 | 0.882 | 0.914 | 0.904 | 0.910 | 0.895 | 0.904 | 0.910 | 0.834 | 0.883 | 0.901 | 0.897 | U2OS.9052.highest.peak.mass |
| 0.945 | NaN | 0.907 | 0.908 | 0.889 | 0.925 | 0.915 | 0.919 | 0.903 | 0.903 | 0.922 | 0.831 | 0.899 | 0.919 | 0.901 | U2OS.9053.highest.peak.mass |
| 0.904 | 0.907 | NaN | 0.909 | 0.874 | 0.911 | 0.882 | 0.882 | 0.874 | 0.888 | 0.897 | 0.810 | 0.863 | 0.882 | 0.890 | U2OS.9508.highest.peak.mass |
| 0.911 | 0.908 | 0.909 | NaN | 0.904 | 0.914 | 0.892 | 0.907 | 0.902 | 0.917 | 0.917 | 0.834 | 0.892 | 0.915 | 0.907 | HCT.9062.highest.peak.mass |
| 0.882 | 0.889 | 0.874 | 0.904 | NaN | 0.877 | 0.867 | 0.896 | 0.897 | 0.901 | 0.881 | 0.853 | 0.889 | 0.903 | 0.871 | HCT.9063.highest.peak.mass |
| 0.914 | 0.925 | 0.911 | 0.914 | 0.877 | NaN | 0.903 | 0.919 | 0.900 | 0.918 | 0.912 | 0.844 | 0.907 | 0.918 | 0.905 | HCT.9064.highest.peak.mass |
| 0.904 | 0.915 | 0.882 | 0.892 | 0.867 | 0.903 | NaN | 0.899 | 0.881 | 0.878 | 0.888 | 0.742 | 0.889 | 0.887 | 0.881 | HeLa.11660.highest.peak.mass |
| 0.910 | 0.919 | 0.882 | 0.907 | 0.896 | 0.919 | 0.899 | NaN | 0.922 | 0.898 | 0.929 | 0.793 | 0.905 | 0.913 | 0.904 | HeLa.12162.highest.peak.mass |
| 0.895 | 0.903 | 0.874 | 0.902 | 0.897 | 0.900 | 0.881 | 0.922 | NaN | 0.881 | 0.894 | 0.749 | 0.900 | 0.896 | 0.873 | HeLa.12166.highest.peak.mass |
| 0.904 | 0.903 | 0.888 | 0.917 | 0.901 | 0.918 | 0.878 | 0.898 | 0.881 | NaN | 0.903 | 0.824 | 0.889 | 0.891 | 0.893 | K562.11661.highest.peak.mass |
| 0.910 | 0.922 | 0.897 | 0.917 | 0.881 | 0.912 | 0.888 | 0.929 | 0.894 | 0.903 | NaN | 0.862 | 0.913 | 0.908 | 0.918 | K562.12159.highest.peak.mass |
| 0.834 | 0.831 | 0.810 | 0.834 | 0.853 | 0.844 | 0.742 | 0.793 | 0.749 | 0.824 | 0.862 | NaN | 0.826 | 0.817 | 0.827 | K562.12160.highest.peak.mass |
| 0.883 | 0.899 | 0.863 | 0.892 | 0.889 | 0.907 | 0.889 | 0.905 | 0.900 | 0.889 | 0.913 | 0.826 | NaN | 0.910 | 0.887 | HEK293.11971.highest.peak.mass |
| 0.901 | 0.919 | 0.882 | 0.915 | 0.903 | 0.918 | 0.887 | 0.913 | 0.896 | 0.891 | 0.908 | 0.817 | 0.910 | NaN | 0.899 | HEK293.11973.highest.peak.mass |
| 0.897 | 0.901 | 0.890 | 0.907 | 0.871 | 0.905 | 0.881 | 0.904 | 0.873 | 0.893 | 0.918 | 0.827 | 0.887 | 0.899 | NaN | HEK293.12019.highest.peak.mass |

**C)** Histograms of highest peak MW values in the range 10-316 kDafor 3 replicates, the average interpolated values and the theoretical MWs (calculated from the sequence) for two representative cell lines, U2OS and HEK293. **D)** Pearson’s correlation coefficients for highest peak MW between all replicates of all cell lines. All data are from Table S2 (10’187 proteins).

Figure S4

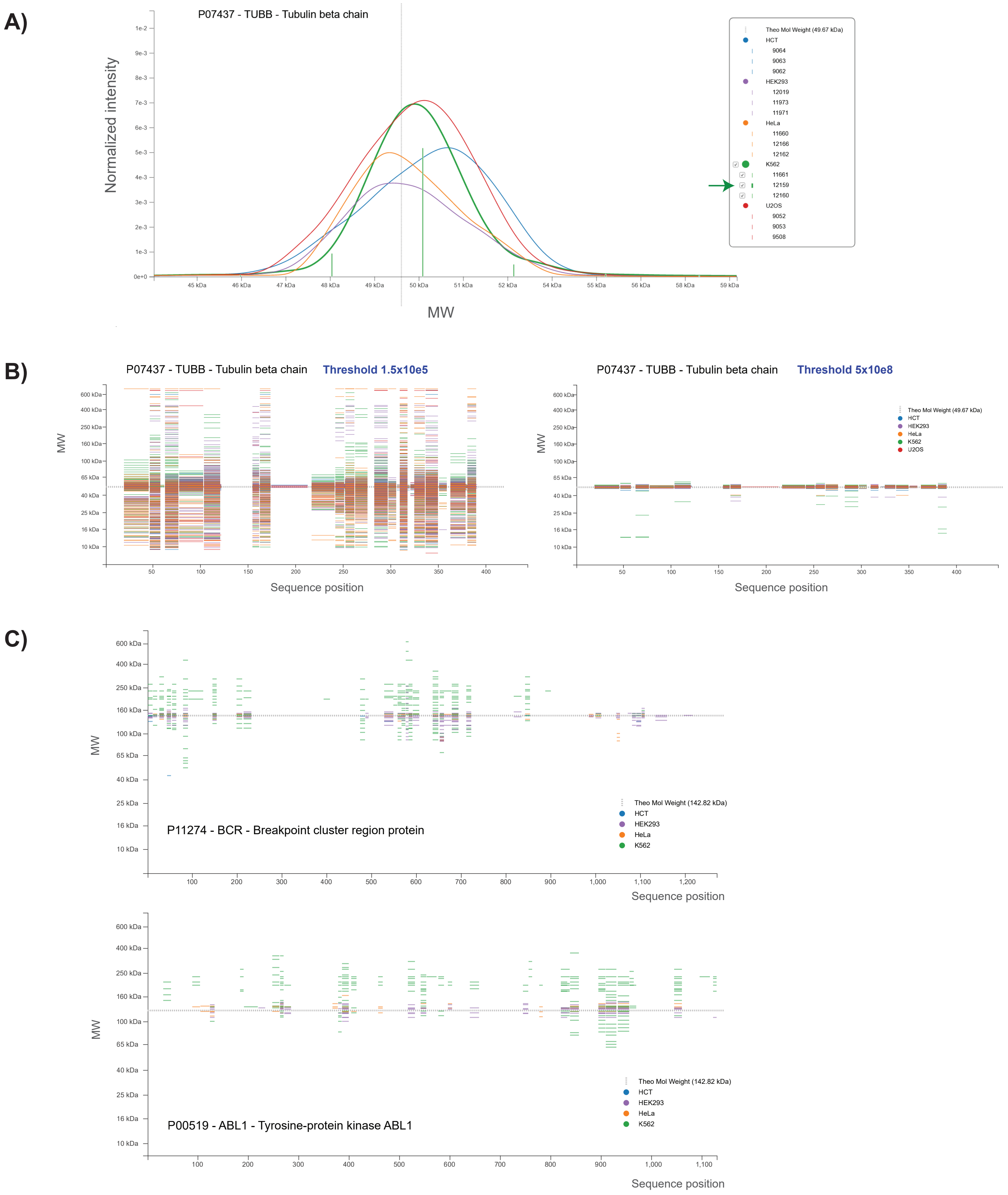


Graphical features and examples of proteins visualized in the PUMBA interface. **A)** detail of graph view for beta tubulin (TUBB). A pop-up menu allows the user to toggle the display of any set of cell lines and replicates. Cursor placement also visualizes raw slice data (vertical bars for K562, replicate 12159). **B)** Intensity threshold adjustments and their impact on the displayed peptide profile for highly abundant proteins such as TUBB that tend to be detected with traces all through the gel. **C)** Peptide profiles for BCR and ABL1 in K562 cells (green) cover the BCR sequence up to residue 901 and the sequence of ABL1 almost entirely, corresponding to the 201 kDa peak observed (Main Fig.4).

Figure S5

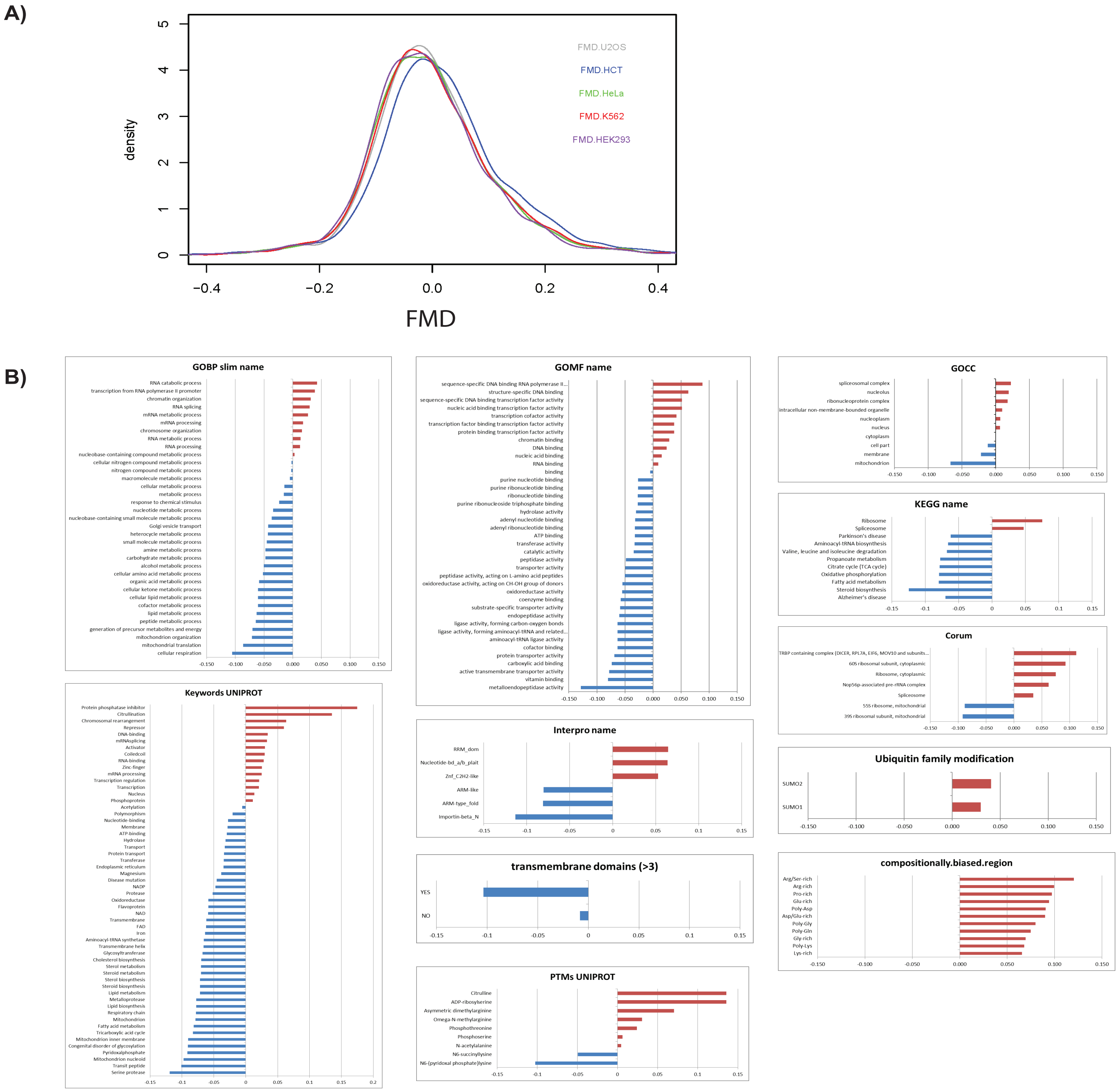


Analysis of Fractional Molecular Weight Deviation (FMD) for 2688 proteins detected in all 5 cell lines with a main single peak accounting for more than 70% total signal. **A)** density plot of FMD values for the 5 cell lines. **B)** Annotation categories whose Fractional Molecular weight Deviation (FMD) were statistically significantly enriched towards either positive or negative values in at least 4 out of 5 human cell lines (1D annotation enrichment ; q-value<0.01; the q-value was the p-value after Benjamini-Hochberg correction). The value on the horizontal axis is the median for each annotation term (blue<0, red >0). With the exception of Gene Ontology terms, all other annotation types were retrieved from the UNIPROT database.

Figure S5 (continued)

C)

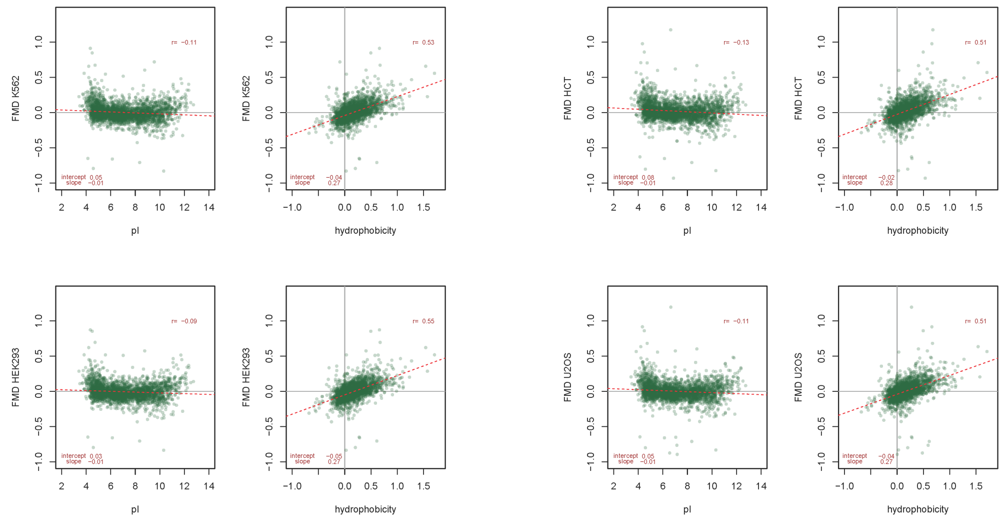


D)

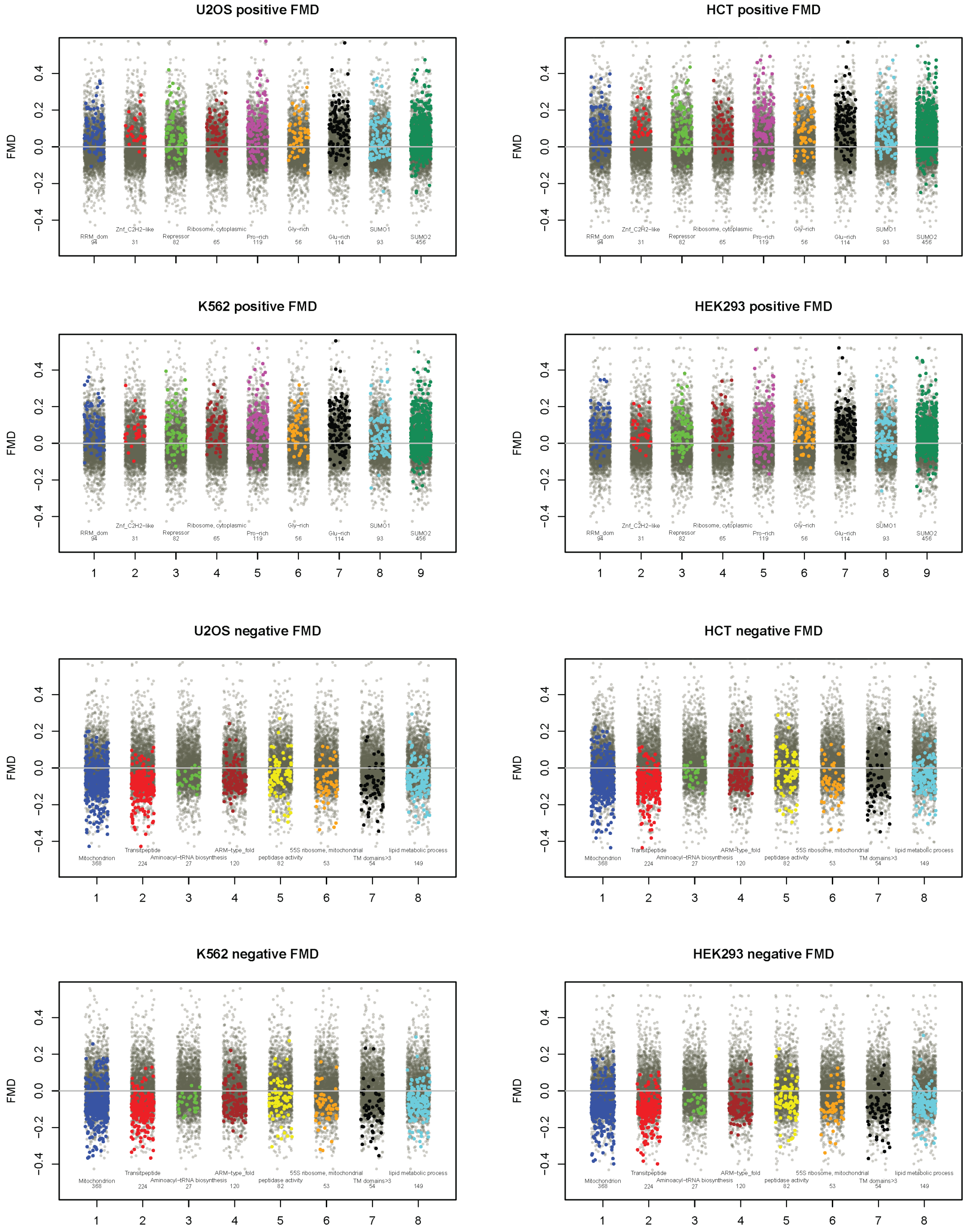


**C),D)** same categories plotted as in main Fig.5, for the other 4 cell lines.
